# A sustained small increase in NOD1 expression promotes ligand-independent oncogenic activity

**DOI:** 10.1101/518886

**Authors:** Leah M. Rommereim, Ajay Suresh Akhade, Bhaskar Dutta, Carolyn Hutcheon, Nicolas W. Lounsbury, Clifford C. Rostomily, Ram Savan, Iain D. C. Fraser, Ronald N. Germain, Naeha Subramanian

## Abstract

Small genetically-determined differences in transcription (eQTLs) are implicated in complex disease but the mechanisms by which small changes in gene expression impact complex disease are unknown. Here we show that a persistent small increase in expression of the innate sensor NOD1 precipitates large cancer-promoting changes in cell state. A ~1.2-1.4 fold increase in NOD1 protein concentration by loss of miR-15b/16 regulation sensitizes cells to ligand-induced inflammation, with an additional slight increase leading to ligand-independent NOD1 activation that is linked to poor prognosis in gastric cancer. Our data show that tight expression regulation of NOD1 prevents this sensor from exceeding a physiological switching checkpoint that promotes persistent inflammation and oncogene expression and reveal the impact of a single small quantitative change in cell state on cancer.

**One Sentence Summary:** A small change in NOD1 expression has a large cancer-promoting impact on cell state.

## Main Text

The innate immune system employs a network of pattern recognition receptors to sense pathogens or danger signals and mount an inflammatory response for host defense. However, chronic low-level stimulation of the immune system can lead to exaggerated responses (*1*) and cause uncontrolled tissue inflammation in genetically predisposed individuals, resulting in autoimmunity, autoinflammatory diseases, and cancers (*2, 3*). Recent studies indicate that in many cases, complex immune-related diseases are linked in genome-wide association studies (GWAS) to causal variants located primarily in non-coding regions of genes, generating eQTLs (expression quantitative trait loci). The extent of expression variation between susceptible and resistant genotypes is often in the 1.5-3 fold range, suggesting that small changes in gene or gene product expression might play an important role in functional immune dysregulation (*4*). While changes in protein concentration or activity due to defects in single genes have been implicated in haploinsufficiencies and monogenic inflammatory or neurodegenerative syndromes (*5*), it is unclear whether a single small alteration in protein concentration can impact the etiology of a multigenic, multifactorial disease like cancer. Recent studies across several cancer types have revealed a regulatory organization in which disease-promoting genomic modifications cluster upstream of functional master regulator proteins whose abnormal activity is necessary and sufficient for propagating a tumor cell state (*6*). Importantly, their activity is controlled post-transcriptionally, and may be dysregulated, for example, by modulation of microRNA (miRNA) activity (*7, 8*) suggesting that small changes in protein expression of master regulators may induce cellular transformation to a cancerous state.

NOD1 is a ubiquitously expressed intracellular innate sensor of microbial infection that senses mesodiaminopimelic acid (iE-DAP, a component of bacterial peptidoglycan) (*9, 10*) and pathogen-induced alterations in cell state (*11*) to trigger nuclear factor-κB (NF-κB) and mitogen-activated protein kinase (MAPK) – dependent induction of pro-inflammatory genes. NOD1 activity is also intimately linked to gastric cancer. Genetic variants in *NOD1* are associated with gastric cancer risk and chronic activation of NOD1 by *Helicobacter pylori* is an initiating event in gastric cancer (*12, 13*). During a study of NOD1 signaling, we discovered that very small changes in NOD1 expression spontaneously lead to large-scale cancer-associated gene activation in monocytes. A big-data approach further revealed that *NOD1* is the most tightly regulated among many innate sensors and suggested a physiological necessity to keep NOD1 expression under stringent control. Here we show that a prolonged small (~1.5 fold) increase in NOD1 concentration within cells upregulates oncogenic activity in the absence of ligand exposure, and identify a miRNA-based circuit for stringent control of NOD1 expression that is relevant in human gastric cancer. Our data highlight one way by which small changes in gene expression can impact the origin of multigenic disease and have broader implications for understanding how small expression changes caused by eQTLs may shape the development of complex diseases like autoimmunity and cancer.

## Results

### NOD1 is activated in a switch-like manner

With an initial aim of investigating the quantitative relationship between NOD1 expression and its effects on cell state, we established a stable, lentiviral expression system in which 3X-FLAG-tagged *NOD1* coding sequence was transcribed under the control of an inducible tetracycline promoter (TRE; tetracycline response element) in THP-1, a human monocyte cell line (THP-1 NOD1 cells) (**Fig. 1A**). As controls, cells transduced with 3X-FLAG NLRP4 (THP-1 NLRP4 cells) or empty lentiviral vector (THP-1 Vector cells) were derived. NLR expression was induced with doxycycline (DOX, an analog of tetracycline) and changes in cellular transcriptome were analyzed by microarray. As expected, addition of DOX led to robust upregulation of *NOD1* and *NLRP4* expression (**Fig. 1B**). The microarray data was further subjected to pathway analysis using GAGE (Generally Applicable Gene-set Enrichment) (*14*), with pathway information derived from KEGG (Kyoto Encyclopedia of Genes and Genomes) (*15*). Multiple pairwise comparisons were conducted to exclude any effects of DOX or the lentiviral expression vector (**Fig. S1**). Surprisingly, this analysis showed that even in the absence of DOX THP-1 NOD1 cells upregulated expression of genes in a large number of cellular pathways similar to that seen upon DOX-induced vast overexpression of *NOD1*, both in the absence of ligand (**Fig. 1C and Table S1**). This behavior was not observed with THP-1 NLRP4 cells or THP1 Vector cells treated + or − DOX suggesting that it was not a general feature of NLRs, DOX treatment, or transduction of cells with the lentiviral vector per se. Careful examination revealed that even in the absence of DOX, there was a ~1.5 fold (1.5X) increase in *NOD1* and ~2X increase in *NLRP4* mRNA in the NLR-transduced cells, as compared to their endogenous levels in the parent or control lentivirus-transduced cells (**Fig. 1B**). Although we did not anticipate this at the conception of the study, this small increase in expression was consistent with previous reports showing that even in the absence of tetracycline the reverse Tet transactivator (rtTA3) binds weakly to the TRE promoter leading to a low-level of background activity (*16*). Quantitative RT-PCR using primers specific for endogenous *NOD1* or 3X-FLAG *NOD1* (**Table S2**) confirmed that THP-1 NOD1 cells showed a small increase in expression of TRE-driven *NOD1* mRNA compared to THP-1 Vector cells in the absence of DOX without affecting endogenous *NOD1* (**Fig. 1D**). Low-level expression of 3X-FLAG NOD1 in the absence of DOX was also observed at the protein level by immunoblot at high exposures and by flow cytometry (**Fig. 1E-F**). Correlation analysis revealed a very high concordance in genes differentially expressed upon low-level (i.e. -DOX) and DOX-induced expression of NOD1 (r^2^=0.9) but not in genes differentially expressed between low-level and DOX-induced expression of NLRP4 (r^2^=0.01) (**Fig. 1G**). These data suggest that a persistent small (1.5X) increase in expression of NOD1 leads to saturating gene activation in a ligand-independent, switch-like manner. A native gel analysis of NOD1 oligomerization showed that persistent 1.5X overexpression of NOD1 induces oligomer formation similar to that seen with ligand (**Fig. S2A**). Moreover, pro-inflammatory genes and negative feedback regulators previously reported to be activated by NOD1 ligand including *IL1B*, *JUN*, *NFKBIA*/IκBα, and *TNFAIP3*/*A20* (*17*) were upregulated in cells with sustained 1.5X increase in NOD1 expression (**Table S3 and Fig. S2B**) indicating that features of the ligand-induced NOD1 response are observed during ligand-independent activation. Importantly, our flow cytometry data (**Fig. 1F**) showed that the small increase in NOD1 expression was distributed across the population suggesting that the gene expression effects were not due to ‘jack-potting’ with very high expression of NOD1 in only a few of the THP-1 cells and emphasizing that it is the very modest 1.5X increase in NOD1 concentration that underlies the response phenomenon. Taken together these data indicate that NOD1 may be activated in a switch-like manner whereby a small increase in its expression mimics full activation.

**Fig. 1.**
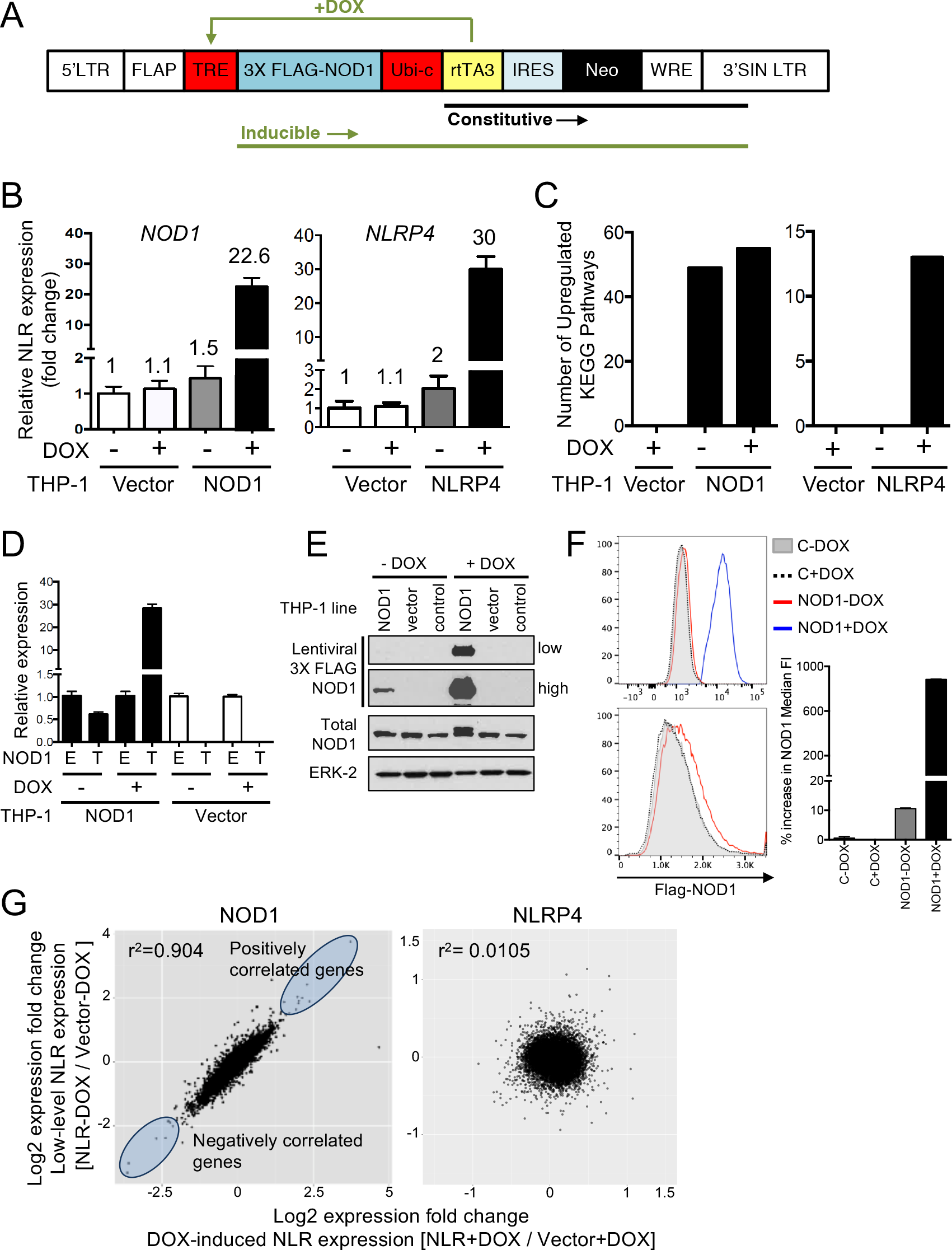
NOD1 exhibits a switch-like behavior. **(A)** Lentiviral system for DOX-inducible expression of NOD1 in THP-1 cells. **(B)** Microarray expression of *NOD1* and *NLRP4* in THP-1 NOD1, NLRP4 and Vector cells treated with (+) or without (−) DOX for 6 h. n=4 per condition. **(C)** Number of KEGG pathways upregulated in the indicated THP-1 cell lines treated as in (B). **(D)** qPCR showing expression of endogenous NOD1 (E) or TRE-driven FLAG-tagged NOD1 (T) in the indicated THP-1 lines treated with or without DOX for 6 h. **(E-F)** Immunoblot (E) and flow cytometry (F) (plots: left; quantification: right) for NOD1 protein in the indicated THP-1 lines with or without DOX. FACS plots show NOD1 expression on a log scale (top) and a linear scale (bottom). **(G)**Correlation plot of genes differentially expressed (fold change>1) in THP-1 NOD1 (left) and THP-1 NLRP4 (right) cells treated as in (B). In (E), ‘control’ refers to THP-1 cells with no lentivirus transduction. Data in D, E and F are representative of at least three independent experiments and the bar graph in F includes pooled data from two independent experiments. Error bars are mean±SEM of four replicates.

The finding that a sustained small change in NOD1 could lead to striking alterations in cellular state prompted a review of gene expression databases to determine how *NOD1* compares to other innate immune sensors in terms of expression regulation. We applied a big data approach to systematically analyze publicly available gene expression data from >70,000 samples from the GEO database corresponding to five widely-used human microarray platforms (GPL96, 97, 571, 5175 and 6480; **Table S4**) and concluded that *NOD1* exhibits the least variation in gene expression among known innate sensors regardless of the experimental context. To make datasets generated in different laboratories cross comparable, genome-wide expression data from each sample was converted to robust z-scores (**Fig. 2A**). For each available probe corresponding to the innate sensors, the variability in distribution of z-score across all the samples was estimated from the standard deviation (SD). From this unbiased and data driven analysis, probes targeting the *NOD1* gene consistently showed minimal deviation in gene expression across all microarray platforms (**Fig. 2B**), implying that *NOD1* expression is under especially stringent control.

**Fig. 2.**
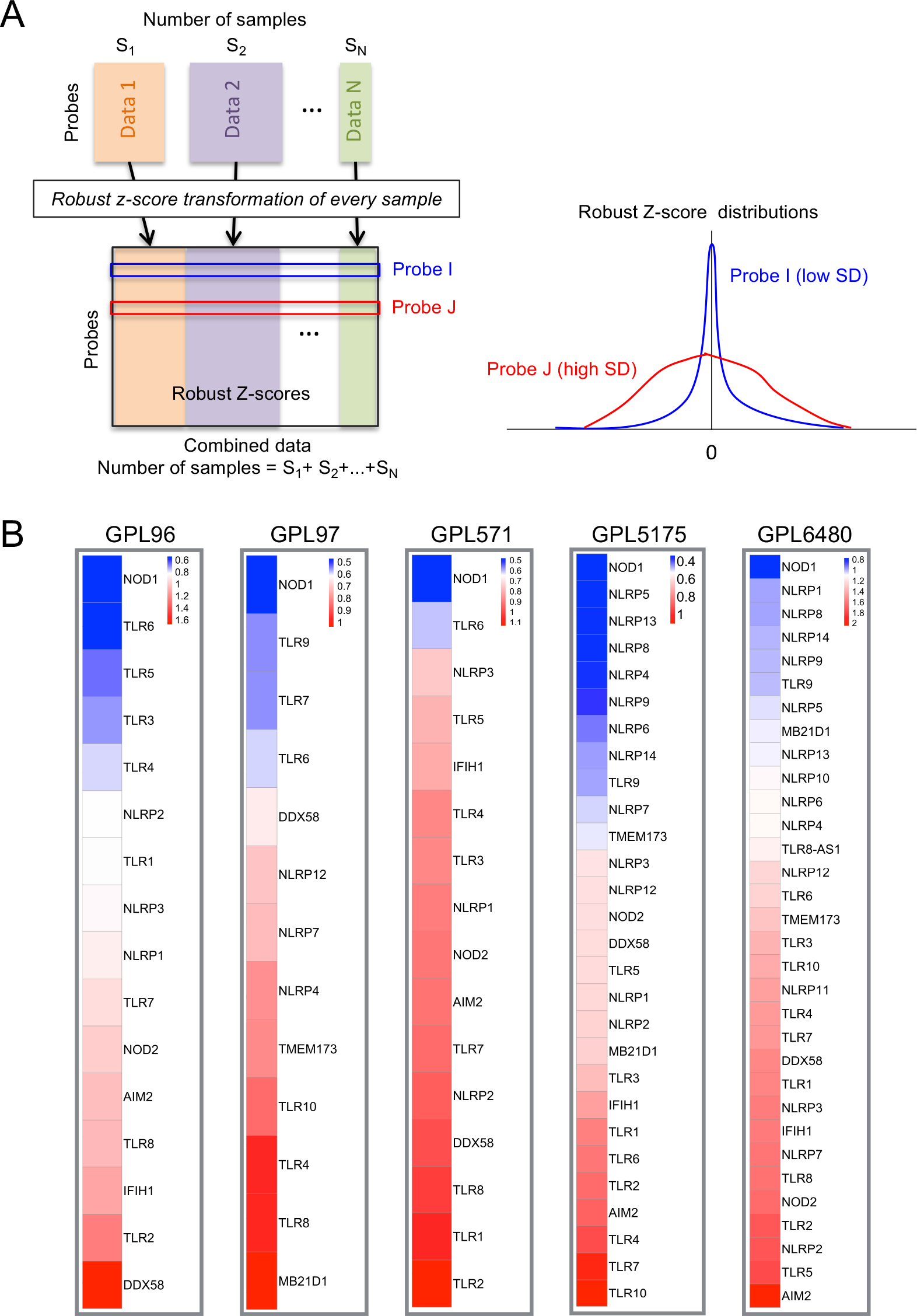
*NOD1* shows minimal change in expression regardless of experimental context. **(A)** Schematic for analysis of publicly available microarray data from the GEO database. Genome-wide expression data from 70,753 samples from five different microarray platforms namely GPL96, 97, 571, 5175 and 6480 were analyzed and transformed into robust z-scores (see methods). For each probe corresponding to innate sensors, the variability in distribution of z-score across all samples was estimated from the standard deviation (SD). **(B)** Heatmaps showing standard deviation in expression of innate sensors from the analyses conducted in A. Genes are arranged in ascending order of standard deviation from top to bottom (blue to red). Not all genes are present on all microarray platforms; for each platform, only genes for which probes were present on that platform are shown.

### MicroRNAs 15b and 16 tightly control NOD1 expression and activation

We next sought to uncover mechanisms of *NOD1* expression control. One class of regulatory elements in the human genome that can exert tight but subtle control of protein expression usually in the less than 2 fold range (*18*) are small noncoding RNAs called miRNAs. The Affymetrix GeneChip Human 1.0 ST microarray used in our study contained probes for 210 pre-miRNAs. Correlation analysis showed that the expression of the pre-miRNAs - mir-15b, mir-16-1 and mir-16-2 - was negatively correlated with *NOD1* but not *NOD2* or *NLRP4* expression (**Fig. 3A and Table S5**). Furthermore, the mature forms of these miRNAs (miR-15b and miR-16) were predicted to bind to the 3’-UTR of *NOD1* in two independent miRNA-target prediction databases - TargetScan and miRanda (**Fig. S3A**; note that the seed regions that mediate complementarity of miR-15b and miR-16 to the *NOD1* 3’-UTR are identical (*19*)). Quantitative RT-PCR showed that expression of miR-15b/16 was reduced to a similar extent in cells with low-level expression of *NOD1* or DOX-induced vast overexpression of *NOD1* when compared to empty vector cells (**Fig. 3B**). Expression of miR-191, an endogenous control, was unchanged. To test if the observed reduction in miRNA under conditions of low-level *NOD1* expression was a result of NOD1 activation, we analyzed expression of miRNAs upon activation of NOD1 with ligand (iE-DAP) in non-transduced, control THP-1 cells. Such activation triggers self-oligomerization of endogenous NOD1 resulting in formation of a protein complex called the nodosome that activates NF-κB and MAP kinases. Indeed, treatment with iE-DAP led to an early decrease in expression of miR-15b/16 by 60 min but this was followed by a restoration of miRNA expression at later times (**Fig. 3C**, **left and middle panels**); this later upswing may be indicative of a ligand-induced negative feedback response. In a reciprocal manner to the miRNAs, the expression of *NOD1* very subtly increased early after ligand stimulation. This was followed by a decline in NOD1 at later times both at the RNA (**Fig. 3C, right panel**) and the protein level (**Fig. 3D**), which coincided with an increase in miR-15b/16 expression. Activation of NF-κB showed similar kinetics and peaked by 60 min after iE-DAP stimulation followed by a decrease by 4 h (**Fig. S3B**).

**Fig. 3.**
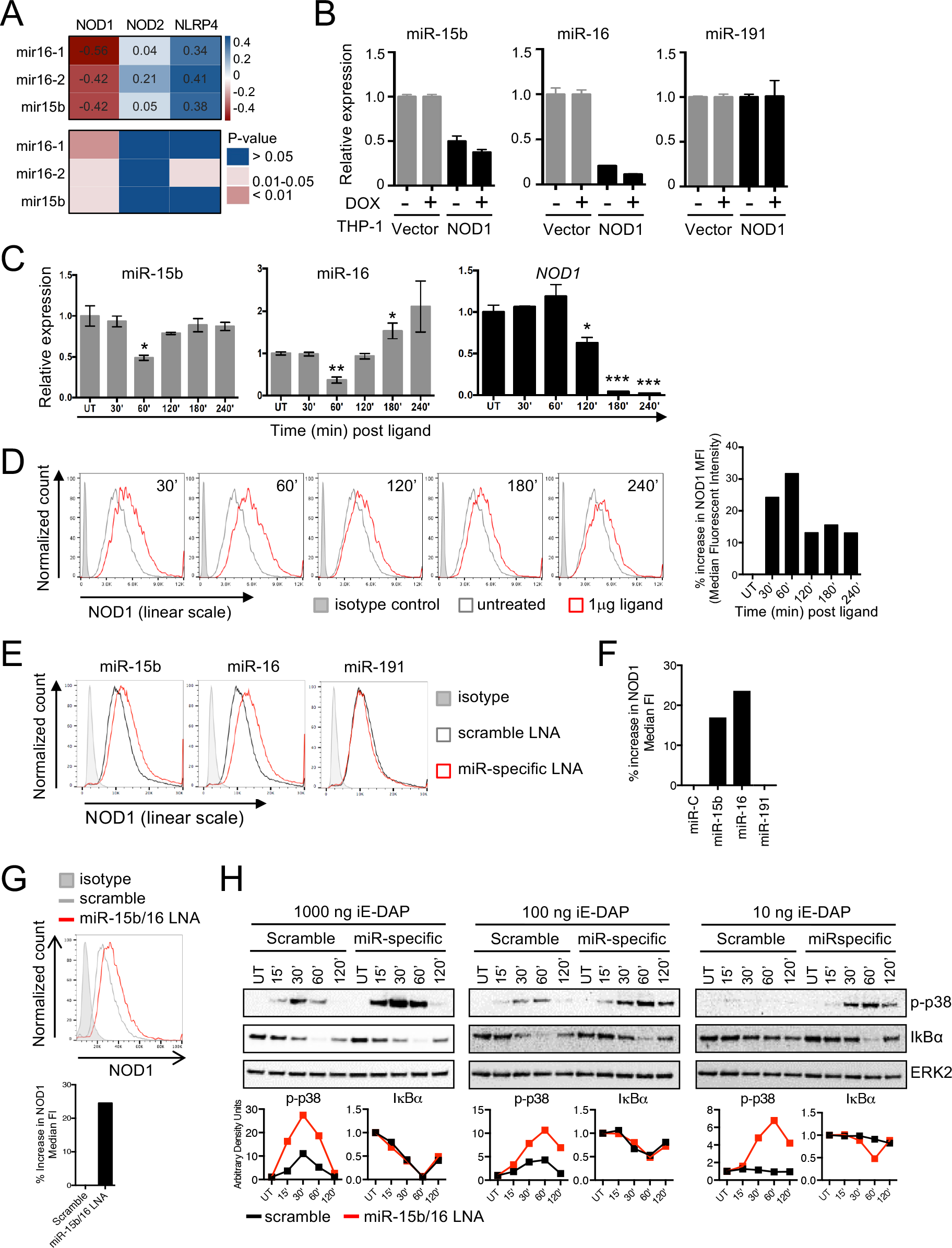
NOD1 expression and activation is tightly regulated by miR-15b and miR-16. **(A)** Top: Correlation (Spearman’s) of mir-15b, mir-16-1 and mir-16-2 with *NOD*, *NOD2* and *NLRP4*. Data originate from quadruplicates of a microarray experiment in THP-1 cells. Numbers indicate r-values and color depicts strength of correlation. Bottom: p-values corresponding to the r-values above. **(B)** q-PCR of miRNA expression in THP-1 NOD1 and Vector cells treated with (+) or without (−) DOX. **(C-D)** qPCR for miRNAs and *NOD1* (C) and FACS for NOD1 (plots, left; quantification, right) (D) in THP-1 cells treated with 1 μg C12-iE-DAP for the indicated times. FACS for NOD1 protein in THP-1 cells transfected with LNA specific to the indicated miRNAs compared to scramble LNA. **(F)** Quantification of histograms in (E). **(G)** Histogram (top) and quantification of NOD1 MFI (bottom) in THP-1 cells transfected with miR-15b/16 LNA compared to scramble LNA. **(H)** Immunoblots (top) and quantification (bottom) of p-p38 and IκBα in THP-1 cells treated with scramble or miR-15b/16 LNA and then left untreated (UT) or treated with the indicated concentrations of C12-iE-DAP for the indicated times. Band intensities of p-p38 and IκBα were normalized to that of ERK2 using ImageJ software. B-H: one representative of at least three independent experiments is shown. Error bars on graphs are mean±SEM of triplicates. * p<0.05, ** p<0.01, *** p<0.001.

Because miR-15b/16 were predicted to bind the 3’-UTR of *NOD1* (**Fig. S3A**), we next examined if these miRNAs in turn control NOD1 expression. To test this, we used highly specific locked nucleic acids (LNA) to achieve short-term inhibition of miR-15b/16 activity in THP-1 cells and observed a subtle (30-40% or 1.3-1.4 fold) increase in NOD1 protein levels in cells transfected with miR-15b/16 LNA compared to cells transfected with control (scramble) LNA (**Fig. 3E-F**). LNA targeting miR-191 had no effect. This phenomenon was also observed in CD14+ monocytes isolated from human blood (**Fig. S3C-D**). Importantly, based on TargetScan and Miranda, binding sites for miR-15b/16 in the *NOD1* 3’-UTR are not conserved between humans and mice. Mouse *Nod1* was not predicted to be targeted by these miRNAs and consistent with this prediction inhibition of miR-15b/16 activity in mouse BMDM did not result in an increase in NOD1 protein levels (**Fig. S3E**) despite abundant expression of these miRNAs in mouse cells at levels similar to those in THP-1 cells (**Fig. S3F**). These data indicate that miR-15b/16 control of NOD1 protein expression is specific to human cells.

We next asked if a small increase in NOD1 by disruption of miRNA control leads to spontaneous NOD1 signaling or sensitizes cells to ligand-induced signaling. To test this, we inhibited miRNA activity in THP-1 cells with LNA and examined NF-κB and MAPK activation in response to iE-DAP. As expected, short-term inhibition of miRNAs led to a small increase in NOD1 expression (**Fig. 3G**) but not a measurable increase in spontaneous NOD1 signaling events. Subsequent treatment with iE-DAP led to enhanced phosphorylation of the MAP kinase p38 in cells treated with miR-15b/16 LNA as compared to those treated with scramble LNA across all ligand concentrations and a delayed degradation of IκBα at a sub-saturating, otherwise inert concentration of ligand (10 ng iE-DAP) (**Fig. 3H**). This is in accordance with a recent work showing that distinct thresholds exist for NF-κB and MAPK signaling downstream of common stimulatory ligand, with the threshold for NF-κB activation being lower than that for MAPK activation (*20*). Taken together, these data indicate that a small increase in NOD1 results in heightened sensitivity of cells to sub-saturating ligand concentrations, a state that might permit commensal products to activate inflammatory responses to a greater degree than when miRNA control is intact.

MicroRNAs can bind the 3’-UTRs of multiple genes. To test if the observed effects of miR15b/16 were directly dependent on their predicted binding site in the *NOD1* 3’-UTR as opposed to indirect effects through binding of these miRNAs to unrelated target genes, we used a target site blocker (TSB), i.e., an antisense oligonucleotide designed to block the miR-15b/16 target site only in the *NOD1* 3’-UTR and not predicted to bind miR-15b/16 sites in any other known human 3’-UTR. As observed with LNA inhibitors, THP-1 cells treated with miR15b/16 TSB showed a subtle (~1.3 fold) increase in NOD1 expression (**Fig. S4A**). This small increase in NOD1 enhanced ligand– induced p38 phosphorylation at all ligand concentrations and IκBα degradation at a suboptimal ligand concentration (**Fig. S4B**), suggesting that miR-15b/16 exert their effect on NOD1 expression and signaling through their predicted binding site in the *NOD1* 3’-UTR.

We next used a luciferase reporter assay system wherein luciferase activity was controlled by the intact *NOD1* 3’-UTR (WT) or a *NOD1* 3’-UTR in which the miR-15b/16 binding site had been mutated (miR-mut) (**Fig. S5A**). Compared to a minimal 3’-UTR (ctrl) the WT *NOD1* 3’-UTR showed reduced luciferase activity when transfected into HEK-293 cells, and mutation of miR-15b/16 binding site rescued this effect (**Fig. S5B**). Co-transfection of the WT and miR-mutant luciferase constructs with miR-15b or miR-16 mimics inhibited luciferase activity of the WT *NOD1* 3’-UTR but had no effect on the miR-15b/16 mutant *NOD1* 3’-UTR (**Fig. S5C**). In contrast, co-transfection of luciferase constructs with LNA to miR-15b or miR-16 increased luciferase activity of the WT *NOD1* 3’-UTR (**Fig. S5D**). Similar results were obtained with the miR-15b/16 TSB, which increased luciferase activity of the WT *NOD1* 3’-UTR but not the miR-15b/16 mutant *NOD1* 3’-UTR (**Fig. S5E**). These results show that miR-15b/16 directly control *NOD1* expression through their specific binding sites in the *NOD1* 3’-UTR.

### A prolonged small increase in NOD1 leads to activation of oncogenes

In contrast to miR-15b/16, which were negatively correlated with *NOD1*, our analyses in **Fig. 1G** revealed that genes whose expression had the greatest positive correlation with *NOD1* expression (r^2^>0.7, fold change≥4) were mainly known proto-oncogenes including *ALX1*, c*-KIT*, *CAV1*, *CNN2*, *CD68* and *GPX8* (**Fig. S6**) (*21–24*). Quantitative RT-PCR, immunoblot and flow cytometry based validation of the two most highly correlated genes ALX1 and c-KIT showed that low-level persistent expression of NOD1 was sufficient to induce maximal expression of these genes, similar to that observed upon DOX-induced upregulation of NOD1 (**Fig. 4A-C**). Like NOD1, the increase in ALX1 and c-KIT was distributed across the population (**Fig. 4C**) indicating that effects on gene expression were not due to very high expression in a few cells, and that it is the very small increase in NOD1 protein concentration that achieves a maximal increase in cancer-related proteins. To determine if a small increase in NOD1 also activates other well-known oncogenes, we analyzed expression of c-MYC and activation (i.e., phosphorylation) of AKT which are considered to be two main drivers in the pathogenesis of many cancers (*25, 26*). Phosphorylation of AKT and expression of c-MYC were both increased in cells with a persistent small increase in NOD1 indicating that unchecked NOD1 signaling upregulates oncogenic processes (**Fig. 4D**). Correlation between expression levels of genes may also imply co-regulation or co-functionality as a result of having common transcription factors (*27*); therefore we next asked if cancer-related genes are coregulated with NOD1. To test this, we stimulated cells with iE-DAP for different times and analyzed expression of *NOD1*, *ALX1* and c-*KIT* by qPCR. The results showed that *NOD1* mRNA subtly increases, and then declines to sub-baseline levels at later times post ligand stimulation, suggesting that *NOD1* expression is likely limited by robust activation-induced negative feedback mechanisms (**Fig. 4E**, **left**). *ALX1* and c-*KIT* mirrored the kinetics of *NOD1* expression (**Fig. 4E**, **middle and right**), suggesting that these genes are co-regulated with *NOD1*. Together these data suggest that *NOD1* and its co-regulated genes are usually under ligand-induced negative feedback that if circumvented by unabated low-level expression of *NOD1*, as seen with the 1.5X over-expression condition (or perhaps with high expression forms of eQTLs) can lead to persistent activation of cancer-associated genes.

**Fig. 4.**
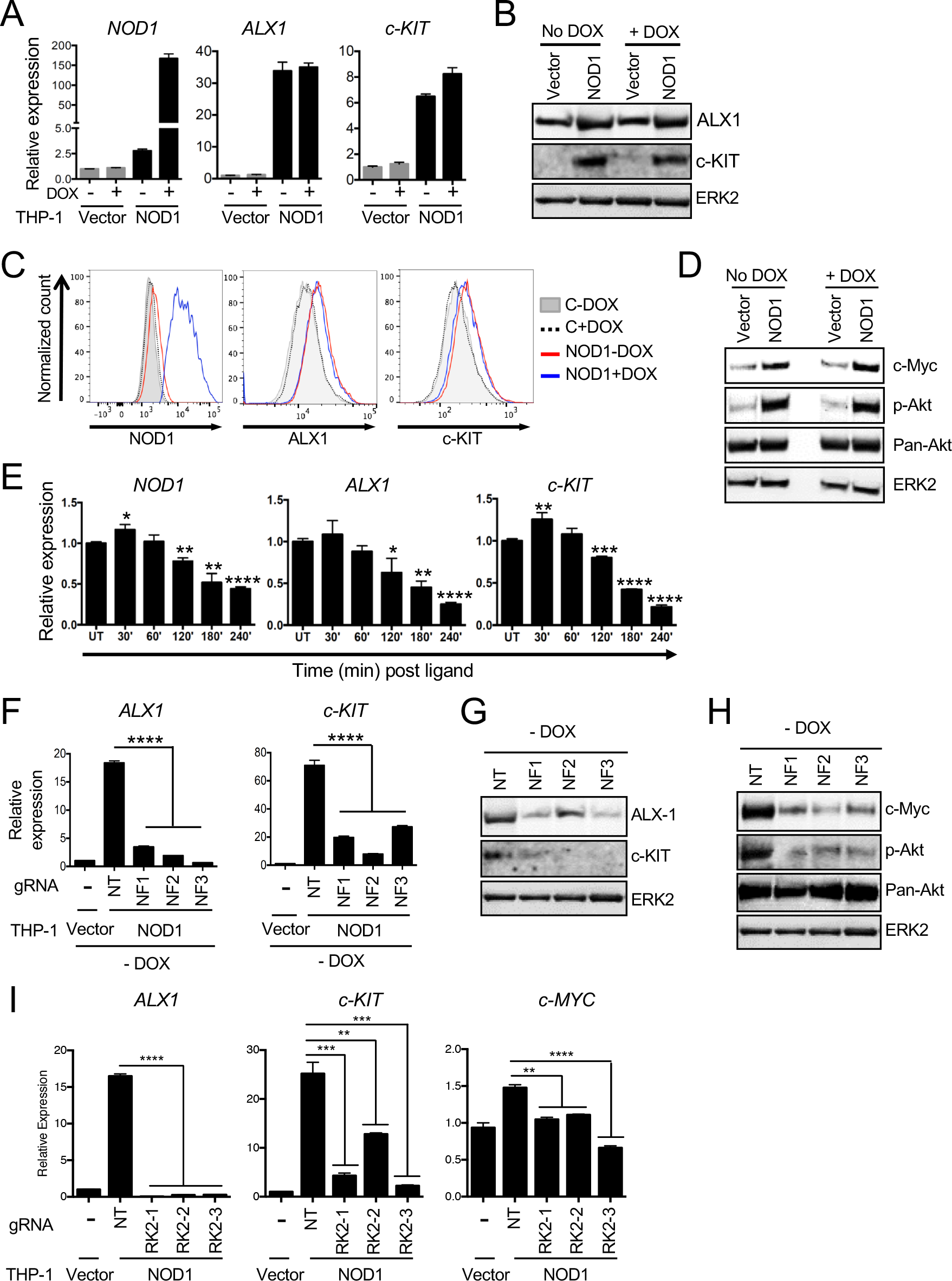
Proto-oncogenes are co-regulated with NOD1 and induced by a sustained small increase in its expression. **(A-C)** qPCR (A), immunoblot (B) and flow cytometry plots (C) showing similarly increased expression of cancer-related genes in THP-1 NOD1 cells in absence and presence of DOX. **(D)** Immunoblot showing similar expression of the oncogenes c-MYC and p-AKT in THP-1 NOD1 cells in presence or absence of DOX. **(E)** qPCR showing expression of NOD1, ALX1 and c-KIT in THP-1 cells treated with 1 μg C12-iE-DAP. **(F-H)** qPCR (F) and immunoblots (G, H) for the indicated genes in THP-1 NOD1 cells following CRISPR/Cas9 mediated ablation of FLAG-*NOD1* in three independent single cell clones (NF1, NF2 and NF3). **(I)** qPCR for the indicated genes in THP-1 NOD1 cells following CRISPR/Cas9 mediated ablation of *RIPK2* in three independent single cell clones (RK2-1, RK2-2 and RK2-3). NT: non-targeting gRNA. NF: gRNA targeting FLAG-NOD1. RK2: gRNA targeting RIPK2. Data are representative of three independent experiments. Error bars on graphs are mean±SEM of triplicates. * p<0.05, ** p<0.01, *** p<0.001, **** p<0.0001.

We next sought to determine if persistent 1.5X over-expression of NOD1 was necessary and sufficient for upregulation of cancer-related genes. To test this, we specifically ablated TRE-driven 3X-FLAG *NOD1* over-expression in THP-1 NOD1 cells using CRISPR-Cas9 based gene editing while leaving expression of endogenous *NOD1* intact and analyzed expression of co-regulated genes. We validated CRISPR targeting in three independent single cell clones by sequencing and observed a unique indel in each clone that resulted in early termination of FLAG-*NOD1* (**Fig. S7A-B**). Ablation of FLAG-NOD1 significantly reduced expression of *ALX1*, c*-KIT*, c-MYC and p-AKT (**Fig. 4F-H**), indicating that the sustained low-level increase in NOD1 expression was causal for upregulation of these genes. Canonically NOD1 signals through its adaptor protein RIPK2. To determine if signaling through this canonical pathway was required for the cancer-related signature observed in response to a sustained 1.5X increase in NOD1 we deleted *RIPK2* in THP-1 NOD1 cells by gene editing and derived three independent single cell clones with unique indels in *RIPK2* that disrupted gene expression (**Fig. S7C-D**). Deletion of RIPK2 reduced the expression of cancer-related genes to near-baseline levels (**Fig. 4I**) indicating that RIPK2 is required for maintaining upregulation of oncogenes in cells with a persistent 1.5X increase in NOD1.

Next, we asked if we could induce a similar oncogenic signature in normal THP-1 cells either by persistently activating NOD1 with ligand or by inducing a prolonged 1.5X increase in expression of endogenous NOD1 through long-term inhibition and/or ablation of miR-15b/16. To test this, we first stimulated THP-1 cells repeatedly with ligand every 24 h for three days, collecting samples every 3-4 h and monitored the expression of *JUN* as a classical ligand-responsive gene and *c-MYC* as a representative oncogene. As expected, ligand treatment led to acute upregulation of *JUN* expression as early as 1 h followed by a gradual tolerization of this inflammatory gene by 48-72 h (**Fig. 5A**). A similar increase followed by a dampening effect was observed with other acute ligand-responsive genes, *IL1B* and *TNFAIP3* (**Fig. S8**); the tolerization effect was also mirrored in cells with persistent 1.5X over-expression of NOD1, which showed lower abundance of these transcripts as compared to that observed at peak ligand stimulation (**Fig. S2B**). In contrast, the expression of c*-MYC* followed reverse kinetics with delayed upregulation between 52-60 h after ligand stimulation which coincided with dampening of the classical ligand-dependent genes (**Fig. 5A**). These data indicate that sustained ligand exposure leads to a delayed shift to oncogene induction. Second, we induced prolonged disruption of miR-15b/16 mediated regulation of NOD1 by treatment of cells with miR-15b/16 TSB that inhibits miRNA binding specifically to the *NOD1* 3’UTR. This led to a small increase in endogenous NOD1 protein by 24 h which was sustained until day 8 of TSB treatment and was accompanied by a significant induction of proto-oncogenes by day 3 (**Fig. 5B-C**). Lastly, we genetically targeted miR-15b and miR-16 in THP-1 cells by CRISPR-Cas9 gene editing in order to achieve a long-term reduction in expression of these miRNAs. miR-15b and miR-16 are present contiguously within an intronic region of the same gene (*SMC4*) and we found that gRNAs targeting miR-15b also significantly reduced miR-16 expression (**Fig. 5D and S9**). Importantly, prolonged reduction in miR-15b/16 expression led to a 1.5 to 2 fold increase in endogenous *NOD1* (**Fig. 5D**) and spontaneous induction of cancer-related genes (**Fig. 5E**). Collectively these data show that a prolonged small increase in NOD1 expression either by transgenic expression from a lentiviral construct or an increase in endogenous NOD1 by disruption of miR-15b/16 function, mimics long-term ligand stimulation and induces expression of proto-oncogenes.

**Fig. 5.**
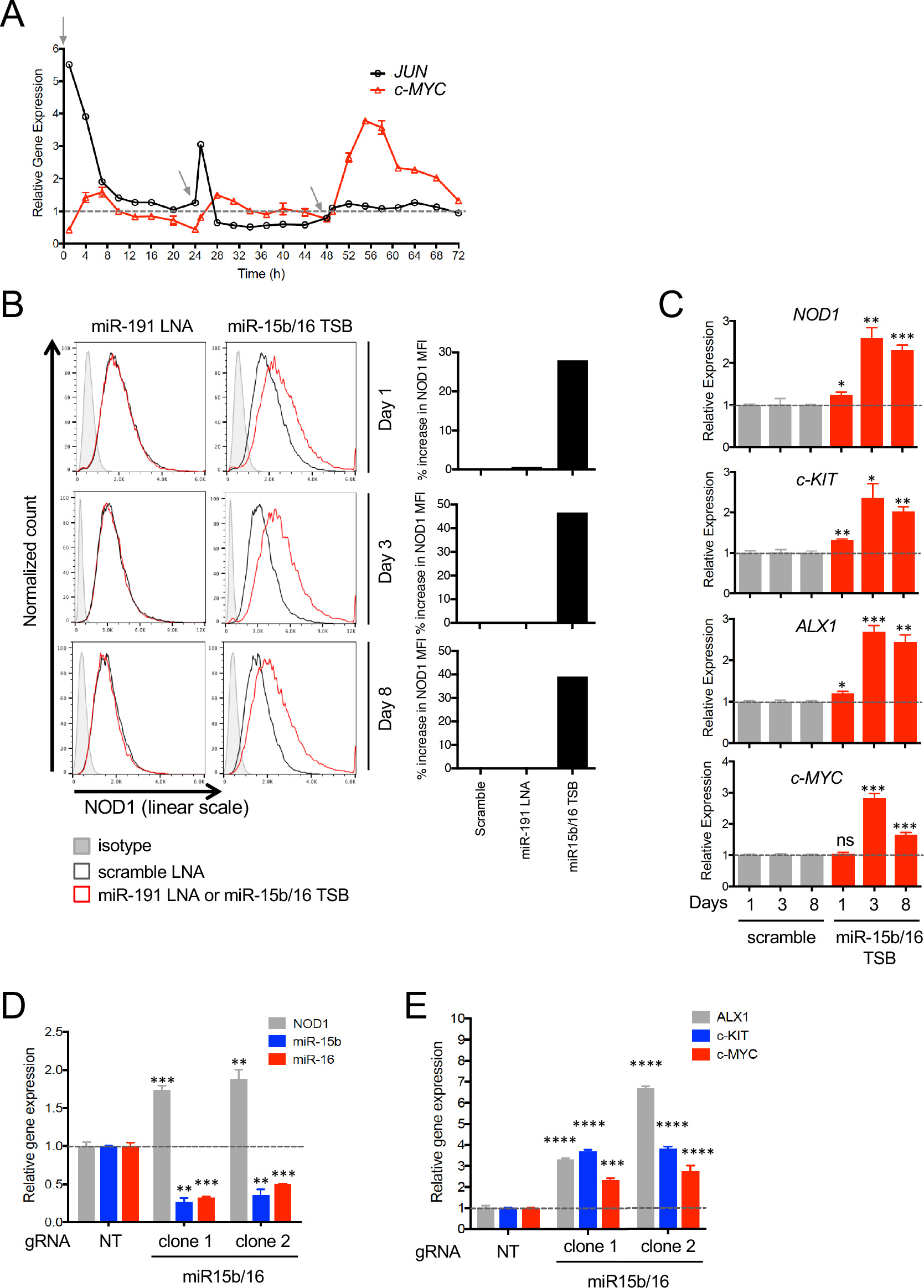
Prolonged activation or increase in expression of endogenous NOD1 induces upregulation of proto-oncogenes. **(A)** Kinetics of *c-JUN* and c-*MYC* expression following ligand treatment as measured by qPCR. Grey arrows indicate times of recurring ligand addition (0, 24 and 48 h). Expression at each time point is represented relative to the untreated control at that timepoint (dashed line set to 1). **(B)** Histograms (left) and quantification (right) showing a small increase in NOD1 protein in THP-1 cells treated with miR-15b/16 Target Site Blocker (TSB). **(C)** qPCR showing increased expression of cancer-related genes in cells treated with miR-15b/16 TSB. Gene expression at each timepoint is represented relative to the scramble LNA-treated control at that timepoint (dashed line set to 1). **(D-E)** qPCR for miR-15b, miR-16 and *NOD1* (D) and the indicated oncogenes (E) in THP-1 cells following CRISPR/Cas9 mediated reduction of miR15b/16 in two independent single cell clones. Gene expression is represented relative to that in the non-targeting gRNA control (dashed line set to 1). Error bars on graphs are mean±SEM and where not visible in panel A are shorter than the height of the symbol.

### A subtle increase in NOD1 predicts poor prognosis in gastric cancer patients

Because a sustained small increase in NOD1 expression was responsible for induction of cancer-related genes, we next asked if dysregulation of miRNA-mediated control of *NOD1* expression is penetrant in human cancer. We first measured correlation between *NOD1* and miR-15b/16 levels in RNA-Seq data from normal vs tumor tissue across 33 different cancer types in The Cancer Genome Atlas (TCGA). This analysis showed that the negative correlation between *NOD1* and miR-15b/16 in normal tissue was most significant in STAD (stomach adenocarcinoma) (**Fig. S10 and Table S6**). This negative correlation was reduced in tumor tissue, suggesting impaired miRNA control of *NOD1* in gastric cancer (**Fig. S10 and S11A**). These findings are consistent with previous reports showing that NOD1 expression is increased in patients with gastritis and gastric cancer (*13*) and expression of miR-15b/16 is reduced in gastric cancer cells (*28*). This relationship between *NOD1* and miR-15b/16 in normal and tumor tissue was not observed in COAD (colorectal adenocarcinoma) (**Fig. S10**) where unlike gastric cancer NOD1 is believed to have a protective role (*29*). Further analysis of RNA-Seq data from the STAD dataset of TCGA (*30*) showed that unlike *NOD1*, negative correlation with miR-15b/16 in healthy gastric tissue and its drop in tumor tissue was not observed for the closely related NLR, *NOD2*, suggesting that regulation by miR-15b/16 is seemingly specific to *NOD1* (**Fig. 6A**). Expression of *NOD1*, but not *NOD2*, was subtly elevated in gastric tumors by ~1.5X when compared to control tissue, an increase consistent with impaired miRNA-mediated regulation of *NOD1* expression (**Fig. 6B-C**). We next analyzed expression of miRNAs and *NOD1* at different stages of the disease, which showed that progression of STAD is associated with increased *NOD1* and decreased miRNA expression (**Fig. 6D and S11B-C**). Taken together, these results imply that reduced expression of miR-15b/16 and a corresponding small increase in *NOD1* expression is linked to lethal gastric cancer progression.

**Fig. 6.**
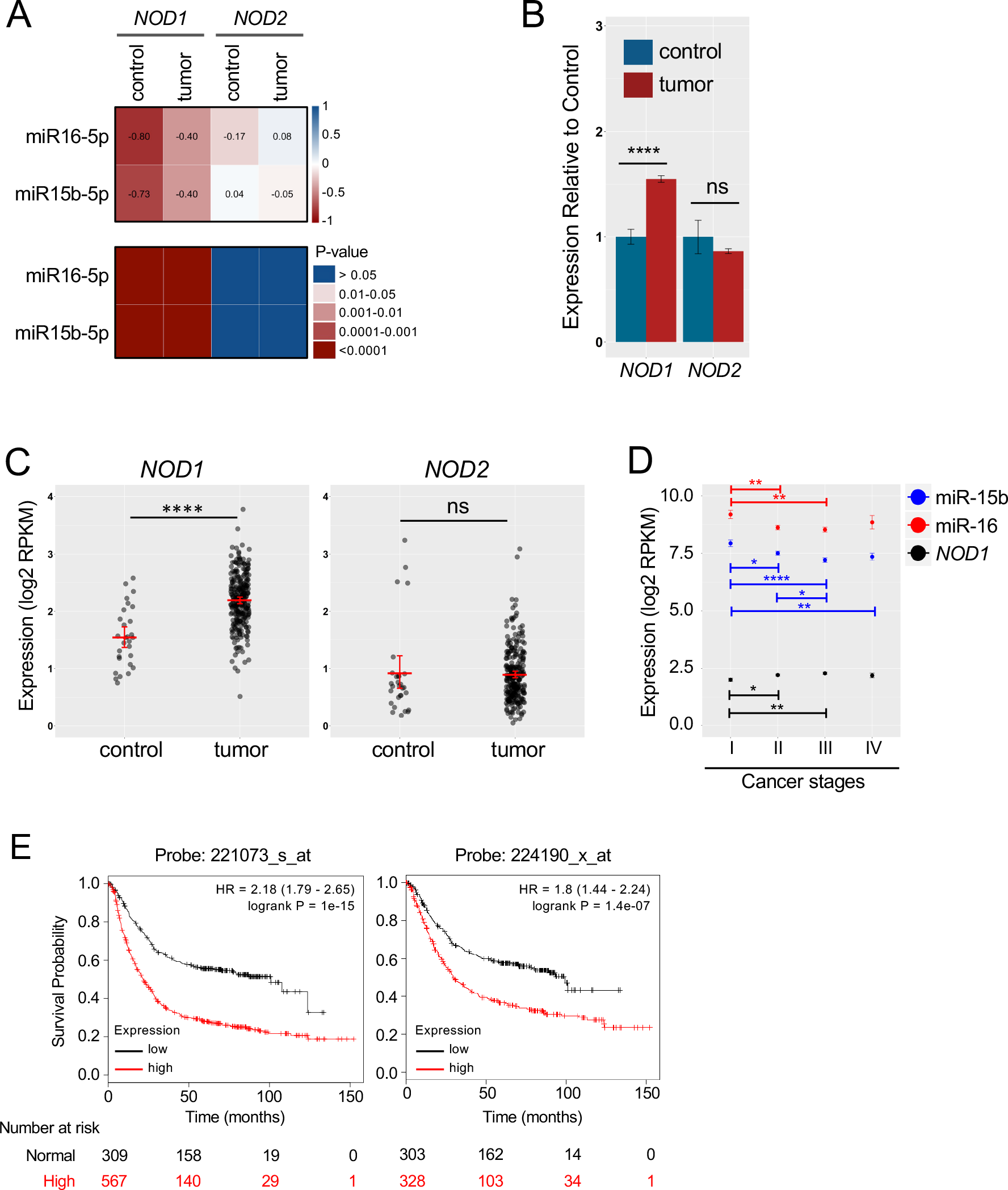
miR-15b/16 regulation of *NOD1* is mirrored in normal human gastric tissue and disrupted in gastric cancer. **(A)** Top: Heatmap showing correlation of *NOD1* expression with miR-15b and miR-16 expression in gastric adenocarcinoma primary tumor tissue (tumor; n=291) and non-cancerous tissue (control; n=29). Numbers indicate r-values and color depicts the strength of the correlation based on r-values. Bottom: Heatmap of p-values corresponding to the r-values above. **(B-C)** Expression of *NOD1* and *NOD2* in gastric adenocarcinoma primary tumor tissue or non-cancerous tissue represented as fold change in expression of each gene in tumor relative to control tissue (B) or absolute log2 RPKM (Reads Per Kilobase per Million) values from RNA-Seq data (C). In (C) red lines denote mean and each dot represents an individual. **(D)** RNA-Seq data showing expression of *NOD1*, miR-15b and miR-16 at different stages of gastric cancer. **(E)** Kaplan-Meier survival curves derived from meta-analysis of survival data from 876 (probe: 221073_s_at; left) or 631 (probe: 224190_x_at; right) gastric cancer patients, showing association of a small increase in NOD1 with patient mortality. B-D: Significance was determined by Welch’s t-test. + denotes significance at p<0.05, ** p<0.01, *** p<0.001, **** p<0.0001, ‘ns’ not significant. Error bars denote SEM from bootstrap.

Dysregulation of miRNA is a well-known feature of cancer. Because a given gene can in principle be targeted by multiple miRNAs, we hypothesized that impaired control of *NOD1* expression in gastric cancer may be a phenomenon regulated by miRNAs beyond miR-15b and miR-16. We therefore expanded our analysis to include all miRNAs that were negatively correlated with *NOD1* expression in normal gastric tissue (r–value between -0.5 and -1) and were also predicted to bind the 3’-UTR of *NOD1*, thereby making them likely candidates to regulate *NOD1* expression. We analyzed correlation of this subset of miRNAs with all NLRs in the TCGA STAD dataset. In healthy gastric tissue, expression of these miRNAs showed the highest negative correlation with *NOD1* compared to all other NLRs (**Fig. S12A-B**). This inverse correlation was reduced in tumor tissue indicating that miRNA control of *NOD1* is globally impaired in gastric tumors. We next asked if a small increase in *NOD1* expression is linked to disease prognosis. To investigate this, we performed meta-analysis of gene expression and survival data from 876 gastric cancer patients (*31*), segregating patients into low and high *NOD1* expressers with a fold difference of ~1.64 between these groups. The results showed that elevated *NOD1* expression is a significant predictor of early disease mortality (**Fig. 6E**). NOD1 and miR-15b/16 may thus serve as biomarkers or predictors of disease prognosis in gastric cancer. Taken together these data imply that miRNAs likely constitute a critical control mechanism that keeps NOD1 expression in check in normal tissue. A small increase in NOD1 is associated with greater disease mortality, underscoring the importance of stringent control mechanisms needed to restrain NOD1 expression.

## Discussion

The human genome contains a large number of regulatory elements that modulate gene activity at different points in the progression from gene transcription to translation. Alterations in non-coding regions of the genome are linked to inflammatory and autoimmune diseases in GWAS (*4*), suggesting that persistent small changes in gene or gene product expression might play an important role in immune dysregulation. Here we provide evidence that a prolonged small increase in expression of NOD1, a ubiquitously expressed cytosolic sensor of bacterial infection, results in a large impact on cell state and dramatic cancer-promoting gene expression in the absence of ligand-driven activity in monocytes. To avoid such spontaneous activity, NOD1 protein expression is very tightly controlled in human cells by at least two miRNAs, miR-15b and miR-16. The fine control of *NOD1* by miRNAs is impaired in gastric tumors, and elevated *NOD1* expression is a significant predictor of mortality in gastric cancer patients (**Fig. S13**). While a direct comparison of the *in vitro* mechanistic data in THP-1 cells or *ex vivo* monocytes from peripheral blood to the gastric cancer data cannot be made due to the heterogeneous nature of normal stomach and gastric cancer tissue, it is worth noting that macrophage infiltration is associated with poor prognosis and increased invasiveness in gastric cancer (*32*) and the gastrointestinal tract is unique in that its macrophages are replenished by blood monocytes (*33*) suggesting that mechanisms observed in monocytes can be impactful in the gastric tumor environment. Future investigations will be required to determine which cells in the heterogeneous samples from gastric tissue have deregulated NOD1.

How might a 1.5X increase in NOD1 expression lead to activation? Because NOD1 needs to oligomerize to signal, we hypothesize that it may operate in a manner akin to a sol-gel transition, whereby a small increase in protein concentration triggers a sharp molecular change from a monomeric species to a macromolecular or polymeric gel-like NOD1 signaling complex that can induce activation of downstream NF-κB and MAPK signaling cascades. Such a phenomenon has been previously observed in multivalent signaling systems including T cell receptor signaling (*34, 35*). Because the protein concentrations required for phase transition depend on physical properties of the monomeric species such as valency and affinity, such a molecular conversion is likely to underlie oligomerization of a large, multivalent entity like NOD1 that has an inherent propensity to not only self-associate but also interact with its downstream signaling adapter protein RIPK2 through homotypic protein-protein interactions. Furthermore, proteins in solution are believed to exhibit dramatic fluctuations in their three-dimensional structures, a movement referred to as protein ‘breathing’ (*36*). Maintaining a supra-physiologic concentration of NOD1 might allow the normal opening and re-closing of each molecule (i.e., molecular breathing) to lead to gel transition that wouldn’t take place when the molecules are more separated and hence, more likely to self-close and cover the oligomerization domain before they associate with another open molecule in the cytosol. Our finding that NOD1 is the most tightly regulated intracellular bacterial sensor (**Fig. 2**) and also that very small increases in intracellular protein concentration drive inflammatory and proto-oncogene expression (**Table S3**, **Figs. 4A-D, 4F-I**, **5B-E**, **S2B and S6**) suggest that evolution has led to NOD1 being maintained at just below the triggering concentration. This yields a highly sensitive detector, at the risk of pathologic activation with small disturbances in expression control.

Recent studies across several cancer types reveal a tumor regulatory architecture wherein functional master regulator proteins whose abnormal activity is necessary and sufficient for implementing a tumor cell state, integrate the effects of multiple and heterogeneous upstream genomic alterations (*6*). In this regard, by virtue of its ability to induce increased expression of cancer-associated genes, NOD1 may itself act as a master regulator whose altered activity propagates a tumor cell state. Recent studies implicate the requirement of at least two signals to transform a normal cell into a tumor cell (*37, 38*). We think it is unlikely that short-term activation of NOD1 alone is sufficient to spur tumor development in otherwise normal stomach tissue because ligand-induced, scaled activation of NOD1 appears to trigger negative feedback mechanisms resulting in repression of NOD1 expression (**Fig. 3C**). Thus, under normal conditions, NOD1 activation is counterbalanced by repression of its expression. To shift this balance towards chronic NOD1 expression it is likely that a disruption of miR-15b/16 and/or other miRNAs that restrict NOD1 expression is required. Importantly, a small increase in NOD1 due to disruption of miR-15b/16 control of its expression brings NOD1 levels closer to the ligand-independent threshold and sensitizes cells to inflammatory responses in response to otherwise inert concentrations of ligand (**Fig. 3G-H and S4A-B**), a state that might permit usually innocuous commensal products to activate inflammatory responses to a greater degree than when miRNA control is intact. Disruption of one or more mechanisms that sustain miRNA expression thus likely constitutes an initial trigger that unleashes the inflammatory potential of NOD1, thereby creating an microenvironment that drives oncogenesis (*3*).

Taken together, our data emphasize that very small prolonged changes in protein concentration can have dramatic effects on cell state that can be penetrant in a disease as complex and multifactorial as cancer. Rather than the ‘Knockout = large effect’ studies that have captivated the field for decades, these data emphasize the need to consider in a more quantitative and subtle way how polygenic diseases may evolve in susceptible hosts over years through a sustained small shift in mean expression of the cells that translates into aberrant population responses over time, with perhaps the occurrence of such events in several genes necessary to escape multilayered control and manifest full-blown clinical disease. More specifically, our findings have important ramifications for focusing on therapeutic approaches that target NOD1 or miR-15b/16 in gastric cancer, and because NOD1 is a ubiquitously expressed mediator of inflammation, may also be of broad relevance to inflammatory cancers that develop beyond the gastric niche.

## Acknowledgments

We thank Dr. Vésteinn Þórsson and Dr. Ilya Shmulevich for helpful discussions on TCGA analysis, Dr. Steve Porcella and the Genomics Unit at Rocky Mountain Laboratories for microarray support, and Dr. Daniel Stetson’s lab at the University of Washington for assistance with gene editing.

## Funding

This work was funded by the Institute for Systems Biology (ISB), U.S. National Institutes of Health grant R21-AI138258, and in part by the Intramural Research Program of NIAID, NIH and funds from the Steven and Alexandra Cohen Foundation to N.S. All work excluding initial acquisition of microarray data and OMiCC analysis was performed at ISB.

## Author contributions

B.D., R.N.G and N.S. conceived the study. R.N.G and N.S. supervised the study and acquired funding. L.M.R, A.S.A. and N.S. designed and performed experiments. L.M.R, A.S.A. and N.S. analyzed the data. B.D., N.W.L., C.H. and C.C.R. performed informatic analysis of microarray, GEO and TCGA data. L.M.R, A.S.A., C.H., C.C.R. and N.S. prepared data for publication. I.D.C.F provided reagents and input for construction of stable cell lines. R.S. advised miRNA target site mutagenesis and luciferase reporter assays. L.M.R. and N.S. wrote the manuscript. R.N.G. and N.S. edited the manuscript. All authors reviewed, revised or commented on the manuscript.

## Competing interests

Bhaskar Dutta is an employee and stockholder of AstraZeneca. None of the other authors have any competing financial interests.

## Data and materials availability

Reagents described in this study are available to the scientific community upon request to N.S. Microarray data generated in this study is available at https://www.ncbi.nlm.nih.gov/geo/query/acc.cgi?acc=GSE99550 and can be accessed for review using the secure token ‘qzajcmwurdcrhkx’. Cancer datasets analyzed in this study are publicly available through The Cancer Genome Atlas and the Kaplan Meier plotter. All other data is available in the main text or the supplementary materials.

