## Supplementary Information for "A sustained small increase in NOD1 expression promotes ligand-independent oncogenic activity"

**This PDF file includes:**

Materials and Methods
Figs. S1 to S13
Tables S1 to S6
References 39-43

### Materials and Methods

Cell lines and reagents

THP-1 cells were obtained from ATCC and cultured in RPMI (Lonza) supplemented with 2 mM glutamine (Gibco), 10 mM HEPES (Corning) and 10% FCS (RPMI-10) in a humidified atmosphere of 5% $CO_2$ at 37°C. Vectors for gateway cloning namely pEN_TmiRc3 entry vector and pSLIK_Neo destination vector were kindly provided by Dr. Iain Fraser, NIAID, NIH. Primers and probes used for detection of endogenous and 3X-FLAG NOD1 and NLRP4 are listed in **Table S2**.

Constructs and stable cell lines

For generation of stable cell lines, 3X-FLAG-tagged NOD1 and NLRP4 were cloned into an entry vector (pEN_TmiRc3) driven by a tetracycline-inducible (TRE) promoter and recombined into a lentiviral expression vector (pSLIK_Neo). Lentiviruses were produced in modified HEK-293T cells (provided by Dr. Iain Fraser) by co-transfecting plasmid DNA of interest along with pRSV, pVSV and pMDL plasmids as previously described (*39*). Virus was concentrated from the supernatant and used to infect THP-1 cells at low copy to ensure <30% infection frequency such that majority of the transduced cells contain a single viral integration. Stable cell lines were selected with 1 mg/ml G418 (Invivogen). Expression of NOD1 and NLRP4 was induced by treatment with 1 µg/ml doxycycline (DOX; Sigma) for 6 h.

Microarray data analysis

THP-1 cells were seeded at a density of 50,000 cells per well in a 96-well plate and treated with 1 µg/ml DOX for 6 hours to induce NOD1 or NLRP4 expression. Total RNA was isolated from 4 replicates per treatment using the RNeasy miniprep kit (Qiagen) and microarray analysis performed using the Affymetrix GeneChip Human Gene 1.0 ST Arrays. Expression data was analyzed by GAGE (Generally Applicable Gene-set Enrichment) (*14*) using the 'gage' package in 'R' (R Core Team (2013). R: A language and environment for statistical computing. R Foundation for Statistical Computing, Vienna, Austria. URL http://www.R-project.org/). Pathway information was derived from KEGG (Kyoto Encyclopedia of Genes and Genomes). Correlations between normalized expression values of miRNAs and *NOD1*, *NOD2* and *NLRP4* were determined using the "rcorr" function with Spearman's method in R.

Big data approach for analysis of expression of innate sensor genes from the GEO database

Genome-wide microarray expression data was downloaded from Gene Expression Omnibus (GEO) to the NIAID high-performance computing cluster by OMics Compendia Commons (OMiCC) project (*40*). Details about quality control, normalization, and annotation are provided in the 'Gene-expression data and pre-processing' section with Supplementary Note 2 of the OMiCC manuscript. Five different human microarray platforms with large numbers of samples (more than 6000 samples per platform) from different vendors were selected for the analysis. These microarray samples were generated from hundreds of microarray experiments. The list of platforms with details on numbers of experiments and samples analyzed from each of these platforms is shown in **Table S4**. For each sample, gene expression values were rescaled by computing a robust z-score. Robust z-scores were used to ensure that outlier expression values, if any, have less significant effect on the rescaled data. Such rescaling of each sample allowed us to compare data across experiments generated by different laboratories. Then, the variability in the distribution of robust z-scores for each probe in a microarray platform across all samples was determined by computing the standard deviation (**Fig. 2A**). These steps were applied for each platform separately and a platform specific standard deviation for each probe was obtained. Finally, probes were mapped to gene symbols using annotation data downloaded from the OMiCC server. All the data and code can be downloaded upon request.

Purification of monocytes from human blood

PBMCs were isolated from heparinized venous blood of healthy adult donors (BenTech) by Ficoll-Paque (GE Healthcare) density gradient centrifugation. CD14+ monocytes were isolated from PBMCs by MACS using Monocyte Isolation Kit II (Miltenyi Biotec). Cell purity was determined by staining with anti-human CD14 (61D3, eBioscience) and analyzed by FACS.

Transfection of cells with LNA inhibitors

MiRCURY LNA$^{TM}$ microRNA power inhibitors were purchased from Exiqon. The following LNA inhibitors were used: hsa-miR-15b-5p, hsa-miR-16-5p, hsa-miR-191-5p, hsa-miR-15b/16 NOD1 3'-UTR specific TSB and scramble negative control A. Briefly, THP-1 cells or *ex vivo* differentiated mouse BMDM were plated at a density of $2.5x10^5$ per well in a 24-well plate in RPMI-10 the day prior to transfection. On the following day, cells were transfected with LNA inhibitors at a final concentration of 50 μM using HiPerfect transfection reagent (Qiagen). Transfection complex was prepared by combining 200 μl OPTI-MEM (Gibco) with 6 μl HiPerfect transfection reagent (Qiagen) and 1μL LNA inhibitor (50 nM stock), and incubated for 20 min at room temperature prior to addition to cells. NOD1 expression was analyzed at 24-36 h post transfection with LNAs. For experiments with ligand, cells were first treated with LNA-based power inhibitors at a final concentration of 75 μM without transfection reagent (to achieve the same inhibition efficacy as that seen with 50 µM inhibitor in presence of transfection reagent), incubated for 24 h and then ligand was transfected using HiPerfect transfection reagent. For long-term experiments with LNA (as in **Fig. 5B-C**), $1.5x10^6$ THP-1 cells were seeded in a 12-well tray, incubated overnight and then treated with 8 µl LNA inhibitor (50 nM stock; without transfection reagent). After 24 h, half of the culture was removed for experiments and an equal amount of fresh media containing LNA (4 µl; 50 nM stock) was added to the well in order to maintain a consistent concentration of cells and LNA inhibitor in each well. This process was repeated for 8 days.

Activation of NOD1 with ligand

THP-1 or CD14+ PBMC cells were seeded at a density of $0.5x10^6$/mL overnight in a 24-well plate in RPMI-10 prior to exposure to C12-iE-DAP (InvivoGen). Ligand was transfected using HiPerfect (Invitrogen). The transfection complex was prepared by combining 200 µl OPTI-MEM with 6 µL HiPerfect and 1 µL ligand (from a 1 mg/mL stock), then incubated for 20 min at RT followed by dropwise addition to each well. Unless otherwise specified, cells were stimulated with 1 µg C12-iE-DAP. At specified times post treatment with ligand, cells were analyzed by flow cytometry, quantitative RT-PCR or cell lysates were prepared and analyzed by immunoblot.

NOD1 3'-UTR Luciferase Reporter Assays

Luciferase reporter constructs were engineered by cloning the WT *NOD1* 3'-UTR or a *NOD1* 3'-UTR with the miR-15b/16 binding site mutated (miR15b/16-mut) (gBLOCK, IDT) into the pGL3 vector (Promega). For luciferase assays, HEK-293 cells were seeded at $4x10^4$ cells/well in a 48-well plate overnight. Cells were transfected with 200 ng NOD1 WT or miR-15b/16-mut 3'-UTR luciferase reporter constructs and 10 ng eGFP using FuGENE HD transfection reagent (Promega).

Where noted cells were co-transfected with either miRNA mimics, LNA inhibitors or TSB at a final concentration of 100 nM using HiPerfect (Invitrogen) for 36-48 h. Cells were washed with PBS and lysed in cell lysis buffer (9803, Cell Signaling Technology). Luciferase activity was measured with the Steady-Glo Luciferase Assay System (Promega) using a microplate reader (Biotek Synergy H4 Plate reader). Luciferase activity was normalized for transfection efficiency (eGFP) and plotted as a fold change over the respective control.

Immunoblot analysis

Immunoblots were prepared using Bolt Bis-Tris gel systems (Invitrogen) and probed overnight at 4°C with antibodies to NOD1 (3545, Cell Signaling), IκBα (4814, Cell Signaling), p-p38 (4511, Cell Signaling), p-Akt (4060, Cell Signaling), pan-Akt (4691, Cell Signaling), c-Myc (13987, Cell Signaling), c-KIT (Ab 81, Santa Cruz), ALX1 (ab181101, abcam) or ERK2 (C-14, Santa Cruz). Anti-FLAG antibody conjugated to HRP (A8592, Sigma) was probed at room temperature for 30 min. Blots were visualized using SuperSignal West Dura chemiluminescent substrate (ThermoFisher) at a high or low exposure as indicated. For native gel analysis of NOD1 oligomers, cells were lysed using lysis buffer containing 1% NP-40. Cell debris were separated by centrifugation at 11000g for 15 min. Native sample buffer was added to the cell lysate and run on a 3-12% native Bis-Tris gel as per the manufacturer's instructions (ThermoFisher). Proteins were transferred to a nitrocellulose membrane followed by immunoblotting.

Quantitative RT-PCR

Total RNA was isolated from THP-1 cells or CD14+ PBMCs using the miRNeasy Mini Kit (QIAGEN) and cDNA was generated using SuperScript III First-Strand Synthesis System

(ThermoFisher). Quantitative RT-PCR was performed using FAM-labeled TaqMan MGB probes (Applied Biosystems) for *NOD1* (hs00196075_m1), *c-KIT* (hs00174029_m1), *ALX1* (hs00232518_m1), *IL1B* (hs01555610_m1), *JUN* (hs01103582_s1), *NFKBIA* (hs00355671_g1), *TNFAIP3* (hs00234713_m1), *GAPDH* (hs02786624_g1), and *ACTB* (hs01060665_g1). Quantitative RT-PCR for hsa-miR-191-5p, hsa-miR-15b-5p and hsa-miR-16-5p was performed using TaqMan MicroRNA Assays (Applied Biosystems). Gene mRNA levels were normalized to the housekeeping genes *GAPDH* or *ACTB* and microRNA levels were normalized to the endogenous control hsa-miR-191-5p.

Analysis of endogenous and transgenic NOD1 expression by flow cytometry

THP-1 or CD14+ PBMC cells were seeded in a 24-well tray at $2.5x10^5$ cells/well in RPMI-10 prior to treatment and were exposed to DOX (1 µg /mL, Sigma) or media for 24 h to induce the expression of NOD1. Alternatively, cells were treated with LNA inhibitors and ligand as described elsewhere in these methods. Endogenous NOD1 expression was assessed using anti-rabbit NOD1 (H-176, Santa Cruz; B-4, Santacruz) and transgenic NOD1 expression was assessed using anti-FLAG M2 antibody (Sigma). Briefly, cells were fixed using CytoFix/CytoPerm (BD Biosciences) followed blocking with Human IgG (Pierce) and staining for primary and secondary antibodies. Cells were analyzed with a BD FACSCalibur flow cytometer (Becton Dickinson). Fluorescence of $10^4$ to $10^5$ cells per sample was acquired in logarithmic mode. Expression was quantified by calculating the mean or median fluorescence intensity of each sample.

Measurement of phospho-p65, ALX1 and c-KIT by flow cytometry

THP-1 or CD14+ PBMC cells were seeded in a 24-well tray at $2.5x10^5$ cells/well in RPMI-10 prior to treatment and were treated with LNA inhibitors and ligand as described elsewhere in these methods. The following antibodies were used: anti-rabbit phospho-p65 (Ser536) conjugated to Alexa Fluor 647 (93H1, Cell Signaling), anti-ALX1 (Sigma, HPA018905) and anti-c-KIT (eBioscience, 17-1178). Briefly, cells were fixed using 4% paraformaldehyde (Alfa Aesar), permeabilized with MeOH followed by blocking with Human IgG (Pierce) and staining with primary and secondary antibodies. After washing, cells were analyzed on a BD FACSCalibur flow cytometer (Becton Dickinson). Fluorescence of $10^4$ to $10^5$ cells per sample was acquired in logarithmic mode. Data was analyzed using FlowJo software.

<u>CRISPR mediated ablation of *RIPK2* and *FLAG NOD1*</u>

For CRISPR/Cas9 targeting of *RIPK2* or FLAG *NOD1*, the guide RNA targeting *RIPK2* or FLAG *NOD1* (IDT) was cloned into a single self-inactivating lentivirus plasmid pRRL-U6-empty-gRNA-MND-Cas9-t2A-Puro, that expresses a Cas9-T2A-puromycin resistance cassette controlled by an MND promoter (*41*). Expression of guide RNA was controlled by the U6-promter. THP-1 NOD1 cells were transduced with lentivirus and stably transduced cells were selected with puromycin (5 μg/ml; Invivogen). Gene targeting events were validated by sequencing and western blot for RIPK2 or FLAG NOD1 to identify $RIPK2^{-/-}$ or $NOD1\text{-}FLAG^{-/-}$ THP-1 cells. The sequence of the gRNA target site is as follows, where a (G) denotes a nucleotide added to enable transcription off the U6 promoter: gRNA targeting *FLAG-NOD1*: (G)AGATCATGATATCGATTACA; gRNA targeting *RIPK2*: (G)AGCGCAGGTCGGCGAGTTTG.

<u>CRISPR targeting of miR-15b/16</u>

Guide RNAs (IDT) were incubated with a fluorescently tagged tracrRNA (IDT) to make a stable duplex, which was then incubated with the Cas9 nuclease (IDT) to make the RNP complex. THP-1 cells were transfected with the RNP complex by nucleofection (Amaxa; SG Cell Line 4-D Nucleofector Kit). Cells were sorted 24 h post transfection by FACS, selecting for cells expressing the tracrRNA fluorescent dye and then subcloned to obtain single cell clones. Gene targeting events were validated by sequencing and quantitative RT-PCR for miR-15b/16. Of the clones reported in this study clone 1 was from a targeting event by miR-15b gRNA #2, clone 2 was from a targeting event by miR-15b gRNA #1. The sequence of the gRNA target site is as follows: gRNA targeting *miR-15b* #1: TGTGCTGTCTACAGTACTTTA; gRNA targeting *miR-15b* #2: AGTACTGTAGCAGCACATC.

TCGA data analysis

Normalized gene expression (RNA-Seq and miRNA-seq) data in gastric adenocarcinoma primary tumor tissue and in non-cancerous tissue was obtained from the gastric adenocarcinoma dataset of The Cancer Genome Atlas (*30*). Analyses were performed in the R environment (*42*), with correlations determined using the "rcorr" function with Pearson's method. For **Fig. S12**, microRNAs predicted to bind to the *NOD1* 3'-UTR were first determined by TargetScan and miRanda. Next, correlations between miRNA and *NOD1* expression in non-cancerous tissue were calculated using the "rcorr" function with Pearson's method in R.  For miRNAs found to be negatively correlated with NOD1 (r  < or = -0.5), correlation was further determined with all NLR genes in both gastric adenocarcinoma primary tumor tissue and in non-cancerous tissue.

Analysis of gastric cancer survival data

We tested association of *NOD1* gene expression levels with survival in gastric cancer patients from a meta-analysis dataset, i.e. a dataset combining the results of multiple independent studies (Affymetrix platforms only). Specifically, we used Kaplan-Meier (KM) Plotter (http://kmplot.com/analysis/index.php?p=background), a web-based tool which maintains a database of gene expression for four different types of cancer, including gastric cancer (*31*). Samples were not restricted based on other factors, such as stage, gender, perforation, treatment, HER2 status. Samples were divided into high and low *NOD1* expression groups and patient survival between these groups was compared. To divide the samples, we used the automatic threshold selection feature of KM Plotter. This functionality carries out KM analysis for all percentiles between lower and upper quartiles and selects the best performing threshold. Two Affymetrix probes were available for the *NOD1* gene in the data sets curated by KM plotter. Details for each of these probes are as follows and survival curves are shown in Fig. 6E.

1. Probe: 221073_s_at; Number of samples: 876; Expression range: 63-1356; Expression threshold: 231; p-value: $1x10^{-15}$; Hazard Ratio: 2.18
2. Probe: 224190_x_at; Number of samples: 631*; Expression range: 4-470; Expression threshold: 98; p-value: $1.4x10^{-7}$; Hazard Ratio: 1.8

* Using the selected probe, only samples measured on HGU133 plus 2.0 arrays can be included in the analysis.

### Quantification and Statistical Analysis

Statistical analyses

Data are presented as means ± SEM (or as indicated in the figure legend). Statistical analyses were performed using Student's t test in either Prism Graphpad or with the "t test" function in R.

p values $< 0.05$ were considered significant. Statistical parameters such as the value of n, the number of replicates, precision measures and statistical significance are reported in the figures and figure legends.

**Supplementary Text**

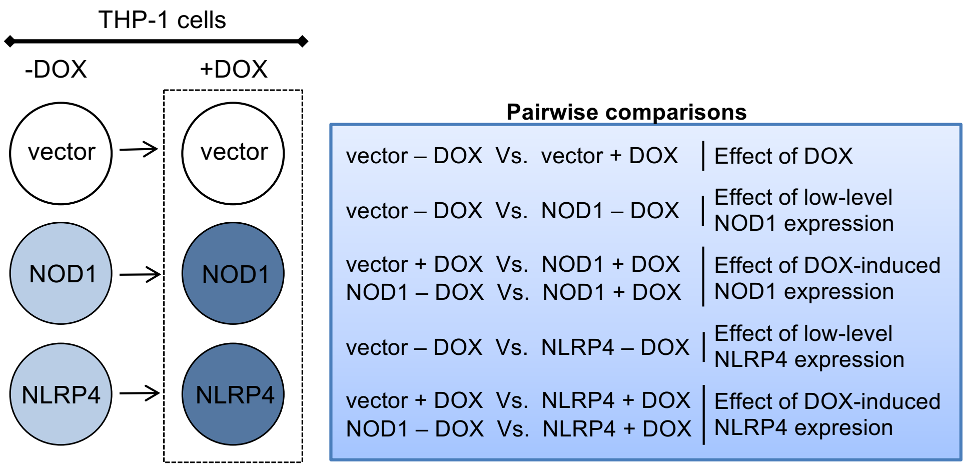


**Fig. S1. Schematic of pairwise comparisons of microarray data.**

Multiple pairwise comparisons conducted for microarray data from THP-1 cells stably transduced with vector alone, NOD1 or NLRP4 are shown. Vector: THP-1 cells transduced with empty lentivirus; NOD1: THP-1 stably transduced with 3X-FLAG NOD1; NLRP4: THP-1 stably transduced with 3X-FLAG NLRP4; DOX: doxycycline.

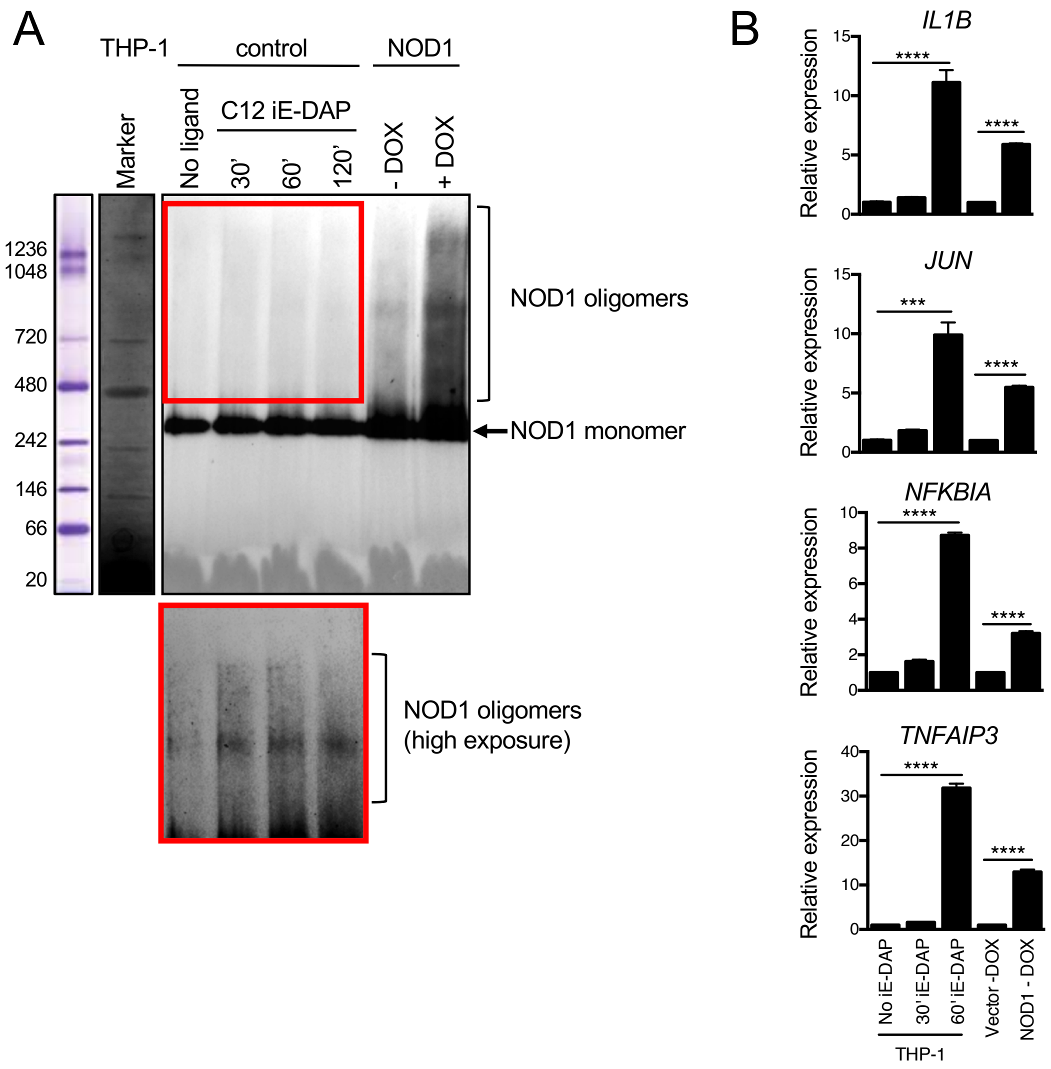


**Fig. S2. A persistent small increase in NOD1 exhibits features of ligand-induced activation and inflammatory gene expression.**

**(A)** Native gel showing NOD1 oligomer formation in cells expressing NOD1 from the lentiviral vector compared to those treated with ligand (10µg C12-iD-DAP for the indicated times). **(B)** qPCR showing that genes conventionally activated by NOD1 ligand are upregulated in cells with a prolonged small increase in NOD1 albeit to a lesser degree. THP-1 cells were untreated or treated with 1 µg C12-iE-DAP for 30 min or 60 min and expression of the indicated genes was analyzed. Data are representative of two independent experiments.

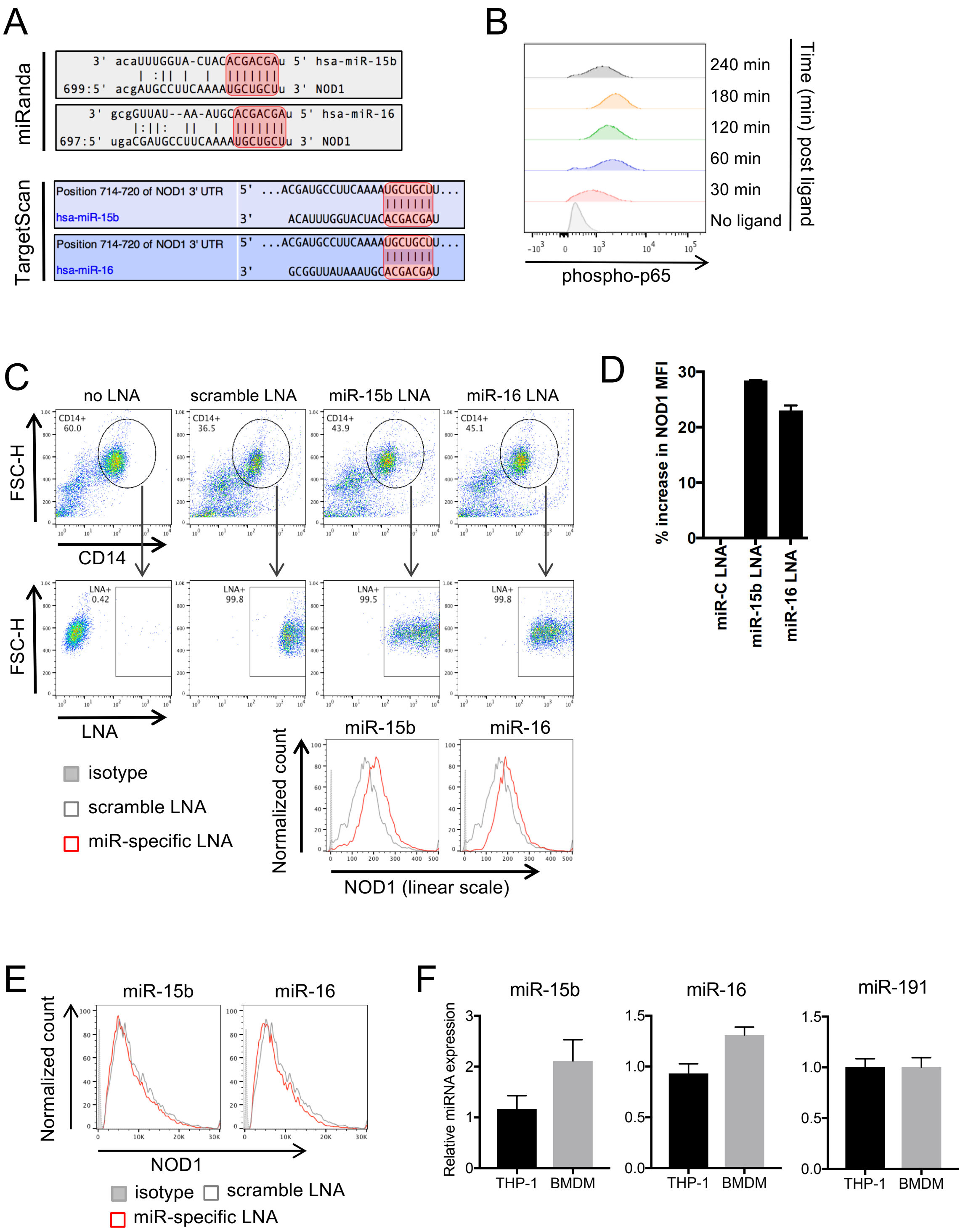


**Fig. S3. miR-15b and miR-16 control NOD1 expression specifically in human cells.**

**(A)** Predicted binding of miR-15b and miR-16 to the 3'-UTR of *NOD1* by TargetScan and miRanda. 7-mer seed regions at the 5' end of the miRNA that exhibit complete complementarity to the *NOD1* 3'-UTR are in red. **(B)** Flow cytometry plots showing phosphorylation of p65 upon C12-iE-DAP treatment. THP-1 cells were left untreated or transfected with 1 μg C12-iE-DAP for the indicated times. **(C)** Representative dot plots (top) showing gating strategy and histograms (bottom) showing expression of NOD1 in CD14+ monocytes transfected with scramble LNA or LNAs targeting miR-15b or miR-16. **(D)** Quantification of NOD1 mean fluorescence intensity (MFI) from two independent flow cytometry experiments. **(E)** Representative histograms from mouse BMDM transfected with scramble or miR-15b/16 LNA showing that expression of mouse NOD1 is not regulated by miR-15b and miR-16. **(F)** qPCR showing relative levels of miR-15b, miR-16 and miR-191 expression in THP-1 cells and mouse BMDM. Note that LNAs targeting miR-15b and miR-16 increase NOD1 expression only in human THP-1 cells (Fig. 3E-F) and have no effect on NOD1 levels in mouse BMDM (Fig. S3E) in spite of similar levels of miRNA expression in these cells.

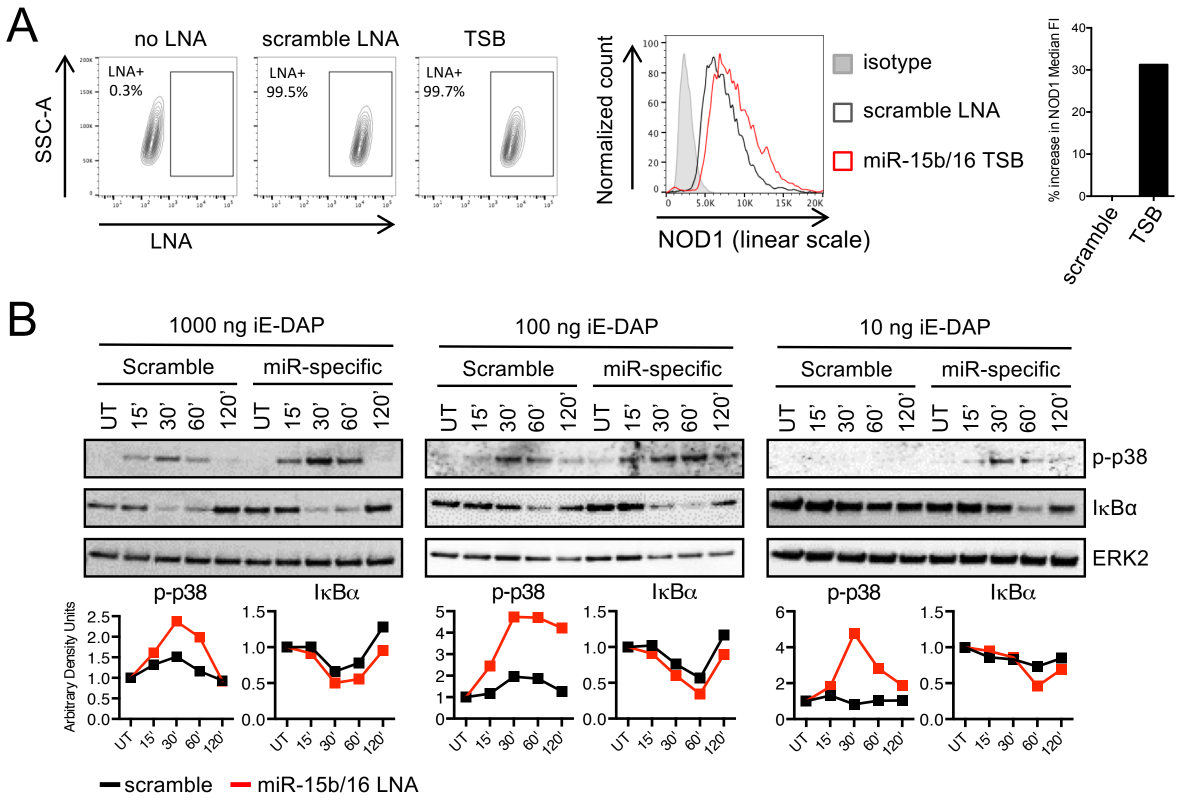


**Fig. S4. Inhibition of miR-15b/16 binding to the *NOD1* 3'-UTR increases NOD1 expression and sensitizes cells to ligand.**

**(A)** Representative FACS plots with gating strategy (left), histogram (center) and corresponding quantification (right) showing subtly increased NOD1 protein in THP-1 cells transfected with Target Site Blocker (TSB) specific for the miR-15b/miR-16 binding site in the NOD1 3'-UTR compared to scramble TSB. **(B)** Immunoblots (top) and corresponding quantification (bottom) of p-p38 and IκBα in THP-1 cells treated with scramble or miR-15b/16-specific TSB and either left untreated (UT) or treated with the indicated concentrations of C12-iE-DAP for the indicated times. Band intensities of p-p38 and IκBα were normalized to that of ERK2 using ImageJ software. Data in A-B are one representative of three independent experiments.

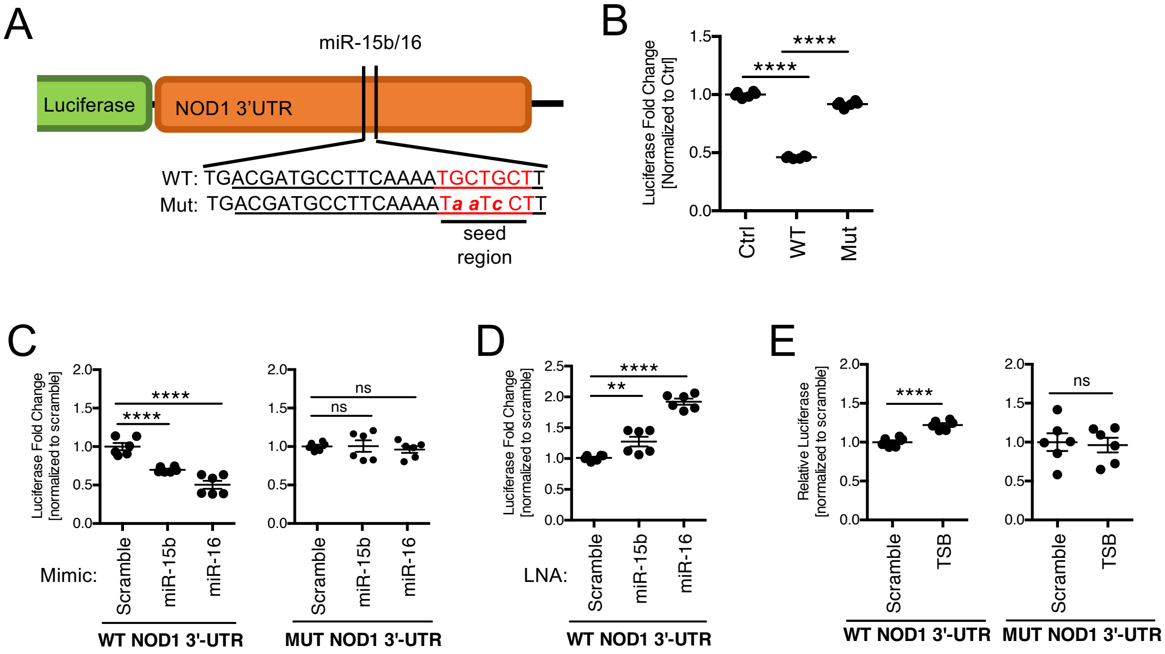


**Fig. S5. miR-15b and miR-16 exert their activity through their binding site in the *NOD1* 3'-UTR.**

**(A)** Schematic of luciferase reporter constructs showing the miR-15b/16 binding site in the *NOD1* 3'-UTR (WT) and mutations introduced to obtain a mutated 3'-UTR (Mut). The seed region is in red and mutations are in lowercase. **(B)** Luciferase reporter assay in HEK-293 cells showing decreased luciferase activity of the WT *NOD1* 3'-UTR compared to a minimal 3'-UTR (ctrl) or a *NOD1* 3'-UTR in which the miR15b/16 binding site was mutated (Mut). **(C)** Luciferase assay showing decreased luciferase activity of the WT *NOD1* 3'-UTR (left) but not the mutated 3'-UTR (right) in response to miR-15b and miR-16 mimics. HEK-293 cells were co-transfected with the indicated 3'-UTR luciferase constructs and miRNA mimics. (**D**) Luciferase assay showing increased luciferase activity of the WT *NOD1* 3'-UTR in response to miR-15b and miR-16 LNA. HEK-293 cells were co-transfected with the WT *NOD1* 3'-UTR luciferase construct and the indicated LNAs. (**E**) Luciferase assay in HEK-293 cells showing increased luciferase activity of the WT *NOD1* 3'-UTR (left) but not the mutated 3'-UTR (right) in response to miR-15b/16 TSB. Data in B-E are representative of at least three independent experiments.

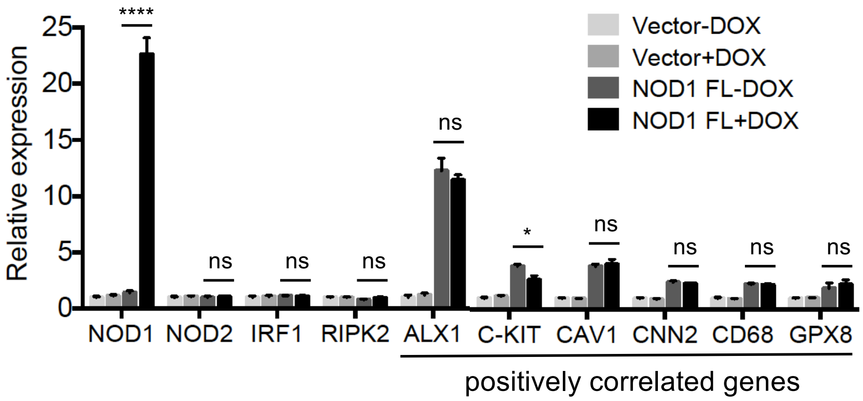


**Fig. S6. A persistent small increase in expression of *NOD1* leads to saturating expression of cancer-related genes.**

Relative expression of the indicated genes from microarray is shown. Error bars are mean±SEM of quadruplicates from a microarray experiment. * p<0.05, ‘ns’ not significant.

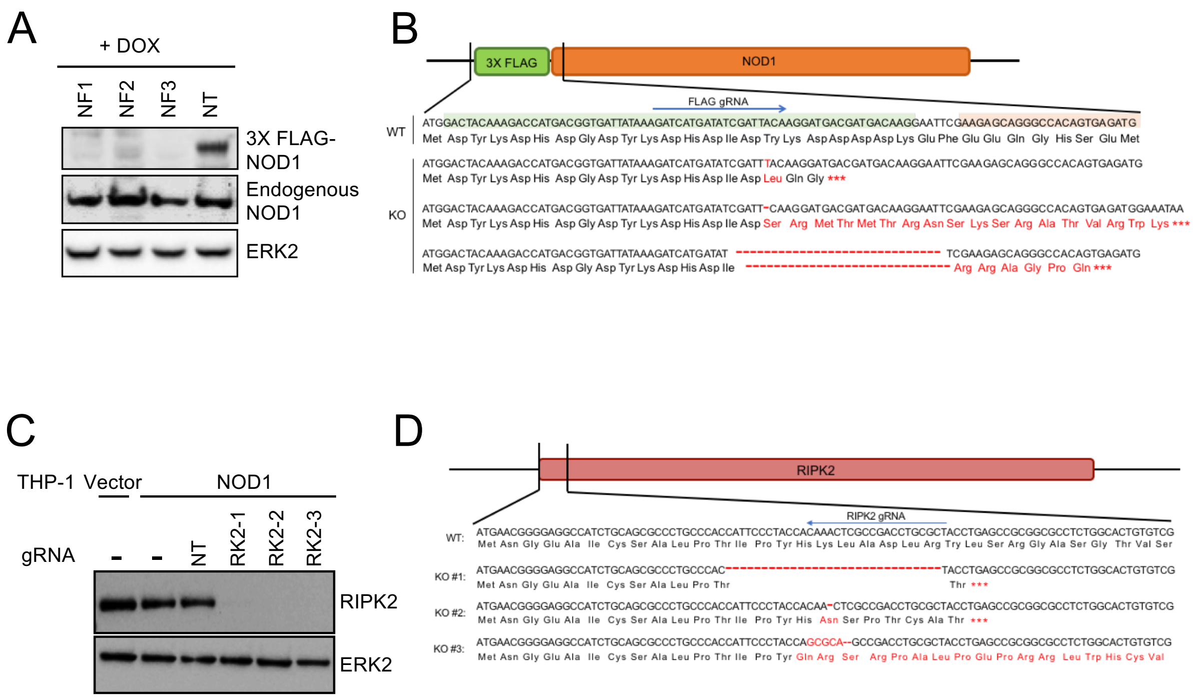


**Fig. S7. CRISPR/Cas9 mediated ablation of FLAG-NOD1 and RIPK2 in THP-1 NOD1 cells.** **(A)** Immunoblot showing the absence of transgenic FLAG-tagged NOD1 protein in THP-1 NOD1 cells following CRISPR/Cas9 mediated ablation of 3X-FLAG NOD1. **(B)** DNA sequencing data showing CRISPR/Cas9 based ablation of FLAG-*NOD1* across three independent single cell clones (KO). **(C)** Immunoblot showing the absence of RIPK2 in THP-1 NOD1 cells following CRISPR/Cas9 mediated ablation of RIPK2. **(D)** DNA sequencing data showing CRISPR/Cas9 based ablation of *RIPK2* across three independent single cell clones (KO). All indels are in red. '----' denotes deleted bases and '***' denotes premature sequence termination at a stop codon. NT: non-targeting gRNA. NF: gRNA targeting FLAG-*NOD1*. RK2: gRNA targeting *RIPK2*.

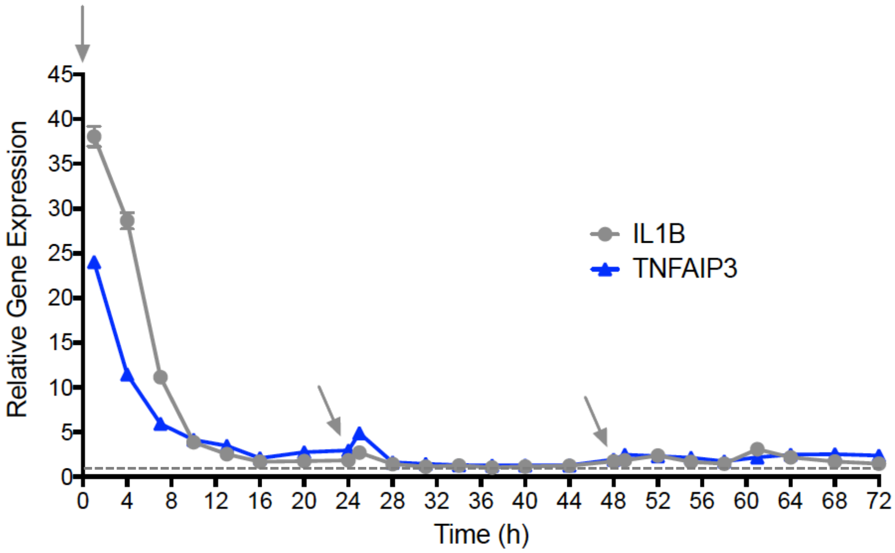


**Fig. S8. Tolerization of acute ligand-responsive genes following repeated NOD1 activation.** Kinetics of *IL1B* and *TNFAIP3* expression following ligand treatment as measured by qPCR. Grey arrows indicate times of recurring ligand addition (0, 24 and 48h). Expression at each time point is represented relative to the untreated control at that timepoint (dashed line set to 1).

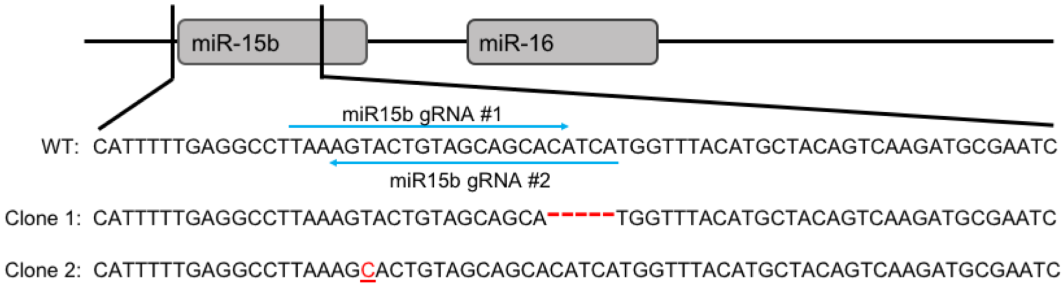


**Fig. S9. Validation of CRISPR/Cas9 targeted editing of miR-15b/16**

DNA sequencing data showing CRISPR/Cas9 based targeting of miR-15b in two independent single cell clones. All indels are in red. '----' denotes deleted bases. Note that miR-15b and miR-16-2 are located contiguously within an intron of the same gene (*SMC4*) and gRNAs targeting miR-15b also reduce expression of miR-16 (Fig. 5D).

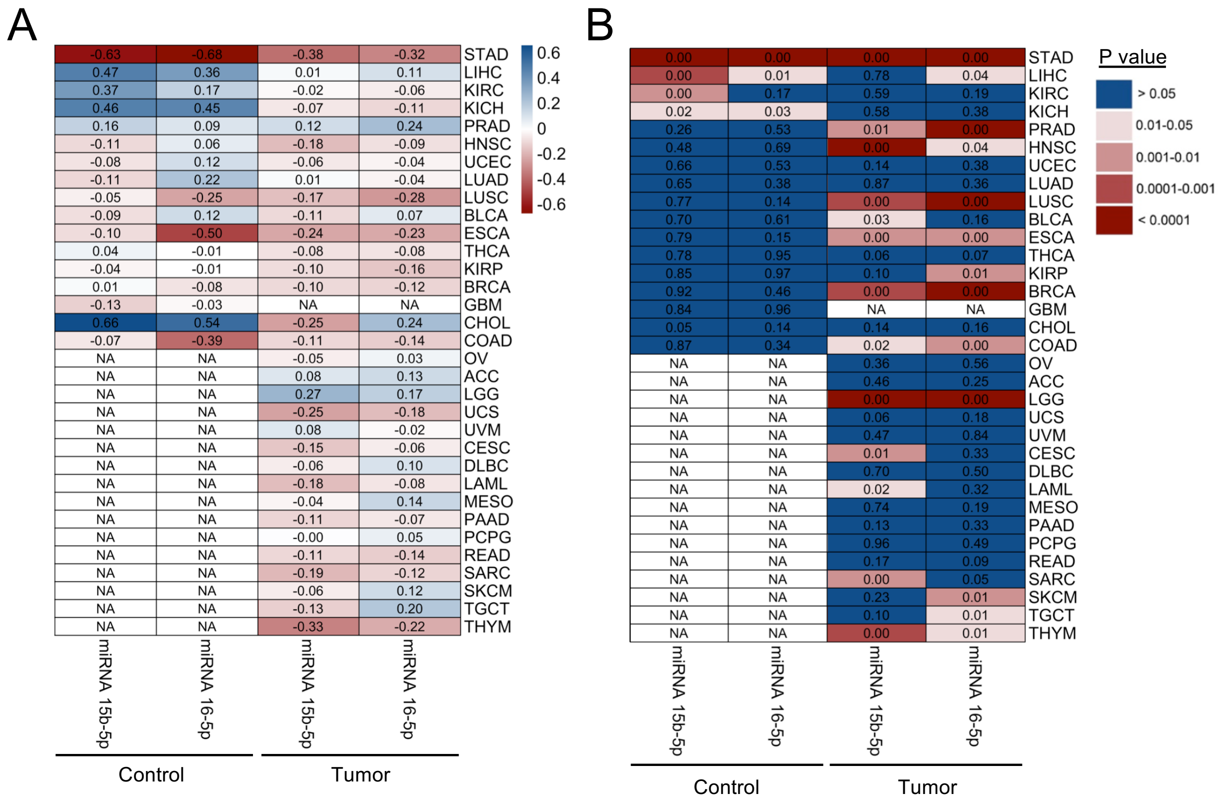


**Fig. S10. Negative correlation between *NOD1* and miR-15b/miR-16 in control tissue and its loss in tumor tissue is most pronounced in stomach adenocarcinoma (STAD).**

(**A**) Heatmap of correlation (r-values) between *NOD1* and miR-15b/16 expression in 33 different cancers from TCGA. r-values are shown for all control and tumor tissues with n>4. NA: n<4. (**B**) Heatmap of p-values corresponding to the r-values in A. Number of samples corresponding to each tumor and control are shown in Table S6.

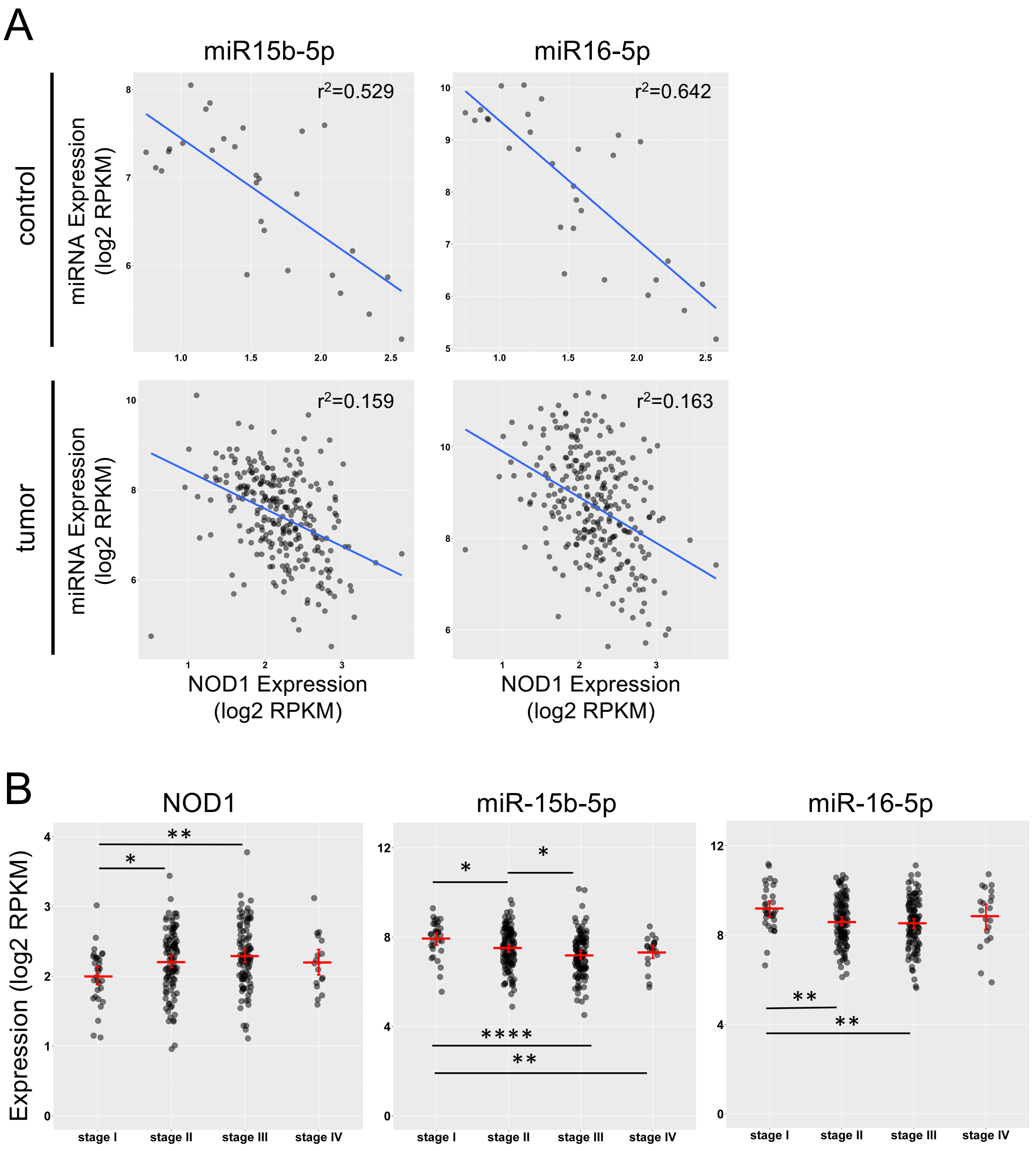


C

NOD1 (p-values)

| | Stage II | Stage III | Stage IV |
|---|---|---|---|
| Stage I | 0.01618 | 0.001463 | 0.1369 |
| Stage II | | 0.2424 | 0.8514 |
| Stage III | | | 0.3823 |

hsa-miR-15b-5p (p-values)

| | Stage II | Stage III | Stage IV |
|---|---|---|---|
| Stage I | 0.01359 | 8.367e-05 | 0.009111 |
| Stage II | | 0.01878 | 0.397 |
| Stage III | | | 0.4351 |

hsa-miR-16-5p (p-values)

| | Stage II | Stage III | Stage IV |
|---|---|---|---|
| Stage I | 0.007481 | 0.002537 | 0.3271 |
| Stage II | | 0.5222 | 0.4681 |
| Stage III | | | 0.3155 |

**Fig. S11. Loss of miR-15b and miR-16 mediated regulation of *NOD1* is associated with progression of gastric adenocarcinoma.**

**(A)** Correlation plots showing decreased negative correlation of *NOD1* with miR-15b and miR-16 expression in gastric adenocarcinoma primary tumor tissue (tumor) compared to non-cancerous tissue (control). $r^2$ values indicate the strength of correlation. Each dot represents an individual. n = 29 for control and 291 for tumor. **(B)** Progression of gastric adenocarcinoma is associated with increased *NOD1* and decreased miRNA expression. Graphs show expression of *NOD1*, miR-15b and miR-16 at different stages of gastric cancer. Data from 291 gastric adenocarcinoma patients sorted by the stage of the disease are shown. Each dot represents an individual. Gene expression was derived from TCGA RNA-Seq data and is shown as log2 Reads Per Kilobase per Million (RPKM). Significance was determined by Welch's t-test. * denotes significance at $p<0.05$, ** $p<0.01$, *** $p<0.001$, **** $p<0.0001$, 'ns' not significant. **(C)** Tables showing exact p-values for comparisons between different stages of gastric cancer corresponding to (B).

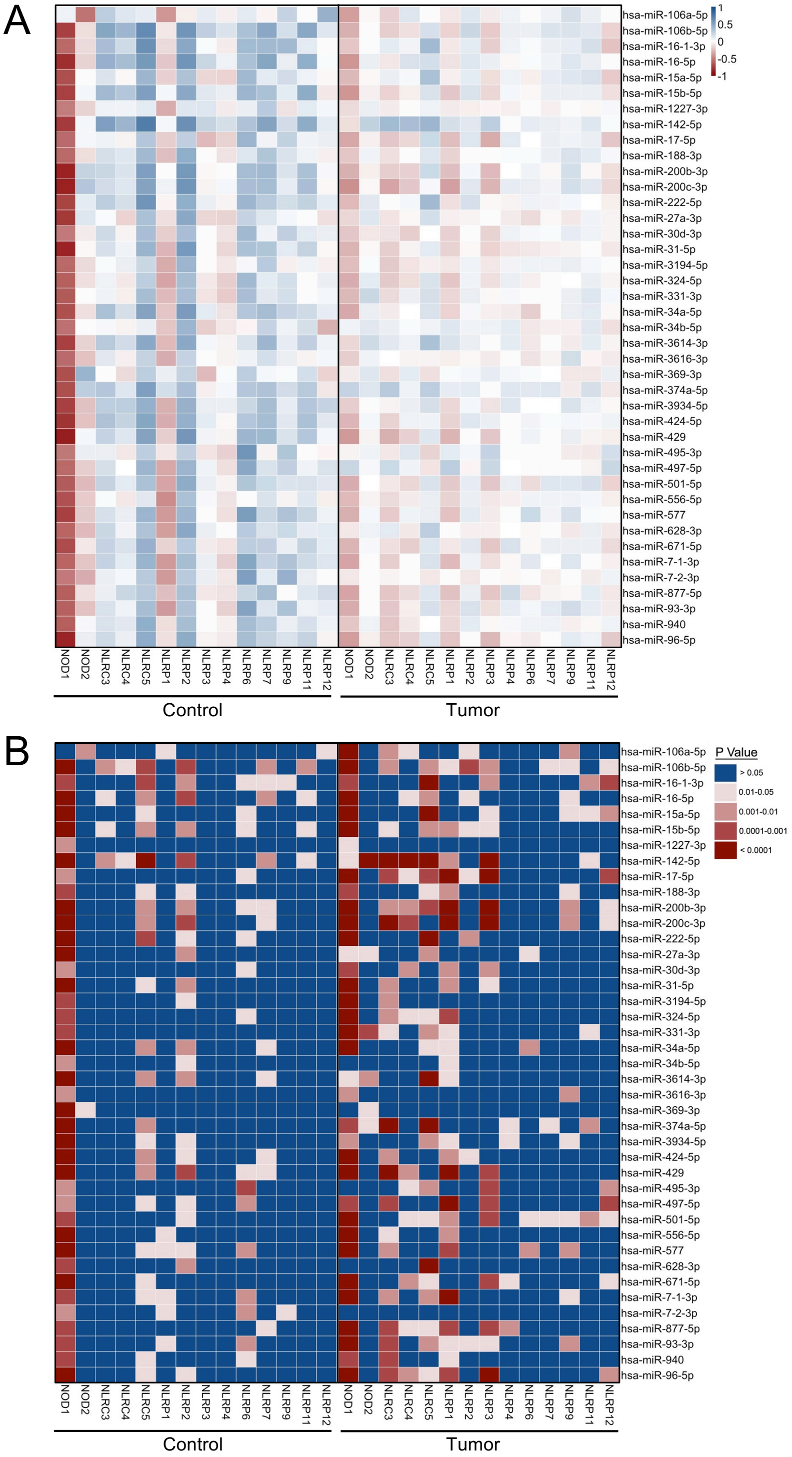


**Fig. S12. Globally impaired miRNA control of *NOD1* in gastric adenocarcinoma.**

**(A)** Heatmap showing correlation (r-values; Pearson's) of expression of NLRs with that of miRNAs predicted to bind the 3'-UTR of *NOD1* from the stomach adenocarcinoma dataset of

TCGA. n = 29 for control and 291 for tumor tissue. **(B)** Heatmap of p-values corresponding to the correlations in A.

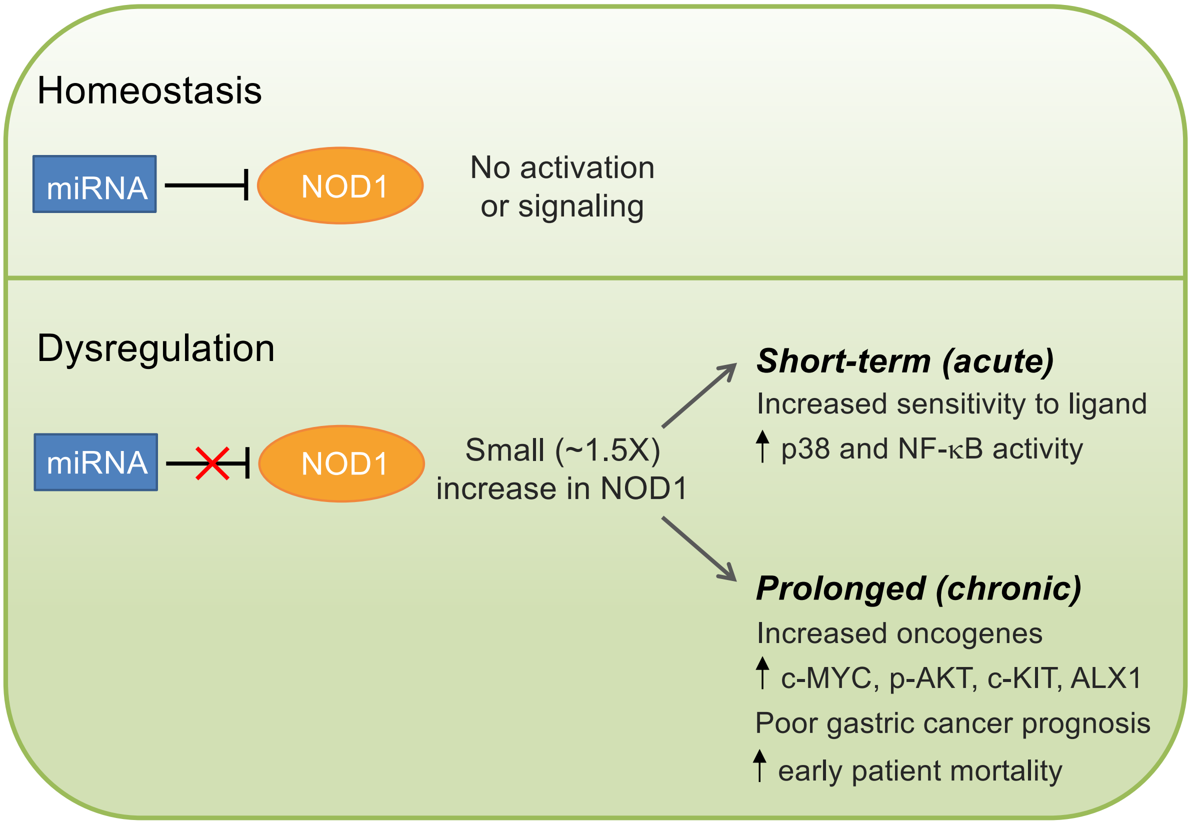


**Fig. S13. Proposed consequences of a small increase in NOD1 expression.**

Under homeostatic conditions, NOD1 expression is kept in check by miRNA (top). Dysregulation of miR-15/16 control of NOD1 (bottom) leads to a small increase in its expression that in the short-term sensitizes cells to inflammatory MAPK and NF-κB signaling in response to sub-saturating, usually inert concentrations of ligand and in the long-term leads to spontaneous activation of cancer-promoting genes. In human gastric cancer, where miR-15b/16 expression is typically reduced or shortening of the 3-'UTR renders NOD1 refractory to regulation by miRNA (*28, 43*), a chronic increase in NOD1 expression and sensitization to MAPK and NF-κB signaling likely creates an inflammatory microenvironment that drives a prolonged switch to oncogenes (*3*).

**Table S1.**

KEGG pathways upregulated by NOD1 and NLRP4 under conditions of low-level (- DOX) or DOX-induced expression.

*Attached as a separate file: Rommereim et al_Table S1*

### Table S2.

| Gene | Oligo Name | Gene Name/Product | 5' – 3' Sequence | Type | Tm | %GC |
|---|---|---|---|---|---|---|
| 3X-FLAG NOD1 transgene | 43-3XFLAGF | FLAG-tag-NOD1 Exon 3 | ATCGATTACAAGGATGACGATGAC | Forward Primer | 58 | 42 |
| | NOD1-30AF126484R | FLAG-tag-NOD1 Exon 3 | GGGTGAGACTCTGATGGGATTATT | Reverse Primer | 58 | 46 |
| | 70-3XFLAG-AF126484FAMRC | FLAG-tag-NOD1 Exon 3 | ACTGTGGCCCTGCTCTTCGAATTCC | 6FAM-BQH1(RC) | 68 | 56 |
| NOD1 endogenous | NOD1-U311NM6092F | Nod1 Exon 2 | GATGGCAAGAGGTGGAGATTG | Forward Primer | 58 | 52 |
| | NOD1-U237NM6092R | Nod1 Exon 2 | TTCCCATAAAAACAGCAACTTGTCT | Reverse Primer | 59 | 36 |
| | NOD1-U263NM6092CFRC | Nod1 Exon 2 | CCCAGATGTTTTCTGTAATCGCCGCC | CalFluorGold540_BHQ1 (RC) | 69 | 54 |
| 3X-FLAG NLRP4 transgene | 55-3XFLAGF | FLAG-tag-NLRP4 exon 1 | GATGACGATGACAAGGAATTCG | Forward Primer | 58 | 45 |
| | 118NM134444R | FLAG-tag-NLRP4 exon 1 | TGAGTTCAAGCTGCAAAGTCATTT | Reverse Primer | 59 | 38 |
| | NLRP4-70NM134444-3XFLAGFAMRC | FLAG-tag-NLRP4 exon 1 | TGAACTCCTCCTTTTTGAGCTCCTCCAGA | 6FAM-BQH1(RC) | 69 | 48 |
| NLRP4 endogenous | NLRP4-U261NM134444F | NLRP4 exon 1 | GGTAGATGAACGCCCTGTGTTT | Forward Primer | 59 | 50 |
| | NLRP4-U186NM134444R | NLRP4 exon 1 | CCTCCACCCCCCATTCTT | Reverse Primer | 58 | 61 |
| | NLRP4-U236NM134444CF | NLRP4 exon 1 | AGGTGCCTCCCAGGAGCCTGAGAC | CalFluorGold540_BHQ1 | 69 | 67 |

List of primers and probes used for evaluating expression of endogenous and 3X-FLAG tagged *NOD1* and *NLRP4* by qPCR.

**Table S3.**

Genes associated with a ligand-induced NOD1 response are upregulated in cells with a small increase in NOD1 but not in cells with a small increase in NLRP4 expression.

*Attached as a separate file: Rommereim et al_Table S3*

**Table S4.**

| GEO Platform ID | Number of samples | Number of experiments | Platform Details |
|---|---|---|---|
| GPL5175 | 8798 | 229 | [HuEx-1_0-st] Affymetrix Human Exon 1.0 ST Array [transcript (gene) version] |
| GPL571 | 10377 | 427 | [HG-U133A_2] Affymetrix Human Genome U133A 2.0 Array |
| GPL6480 | 10930 | 473 | Agilent-014850 Whole Human Genome Microarray 4x44K G4112F (Probe Name version) |
| GPL96 | 34562 | 978 | [HG-U133A] Affymetrix Human Genome U133A Array |
| GPL97 | 6086 | 142 | [HG-U133B] Affymetrix Human Genome U133B Array |

Details of genome-wide microarray expression datasets analyzed in Fig. 2.

**Table S5.**

List of miRNAs on the Affymetrix GeneChip Human 1.0 ST microarray. Correlation of miRNAs with *NOD1* (r-values) and predicted binding of miRNAs to the *NOD1* 3'-UTR are shown.

*Attached as a separate file: Rommereim et al_Table S5*

**Table S6.**

| Cancer type | # Controls | # Patients |
|---|---|---|
| STAD | 35 | 409 |
| LIHC | 49 | 367 |
| KIRC | 70 | 499 |
| KICH | 24 | 65 |
| PRAD | 52 | 491 |
| HNSC | 44 | 513 |
| UCEC | 32 | 522 |
| LUAD | 19 | 507 |
| LUSC | 37 | 464 |
| BLCA | 19 | 405 |
| ESCA | 10 | 182 |
| THCA | 59 | 508 |
| KIRP | 32 | 286 |
| BRCA | 90 | 1073 |
| GBM | 5 | 0 |
| CHOL | 9 | 36 |
| COAD | 8 | 423 |
| OV | 0 | 305 |
| ACC | 0 | 78 |
| LGG | 0 | 528 |
| UCS | 0 | 56 |
| UVM | 0 | 80 |
| CESC | 3 | 305 |
| DLBC | 0 | 47 |
| LAML | 0 | 168 |
| MESO | 0 | 87 |
| PAAD | 4 | 178 |
| PCPG | 3 | 183 |
| READ | 3 | 152 |
| SARC | 0 | 258 |
| SKCM | 1 | 449 |
| TGCT | 0 | 155 |
| THYM | 2 | 120 |

Number of samples corresponding to each tumor and control tissue in Figure S10.
